# TBPL2/TFIIA complex establishes the maternal transcriptome by an oocyte-specific promoter usage

**DOI:** 10.1101/2020.06.08.118984

**Authors:** Changwei Yu, Nevena Cvetesic, Vincent Hisler, Kapil Gupta, Tao Ye, Emese Gazdag, Luc Negroni, Petra Hajkova, Imre Berger, Boris Lenhard, Ferenc Müller, Stéphane D. Vincent, László Tora

## Abstract

During oocyte growth, transcription is required to create RNA and protein reserves to achieve maternal competence. During this period, the general transcription factor TATA binding protein (TBP) is replaced by its paralogue, TBPL2 (TBP2 or TRF3), which is essential for RNA polymerase II transcription. We show that in oocytes TBPL2 does not assemble into a canonical TFIID complex. Our transcript analyses demonstrate that TBPL2 mediates transcription of oocyte-expressed genes, including mRNA survey genes, as well as specific endogenous retroviral elements. Transcription start site (TSS) mapping indicates that TBPL2 has a strong preference for TATA-like motif in core promoters driving sharp TSS selection, in contrast with canonical TBP/TFIID-driven TATA-less promoters that have broader TSS architecture. Thus, we show a role for the TBPL2/TFIIA complex in the establishment of the oocyte transcriptome by using a specific TSS recognition code.

## Introduction

Regulation of transcription initiation by RNA Polymerase II (Pol II) is central to all developmental processes. Pol II transcription requires the stepwise assembly of multi-protein complexes called general transcription factors (GTFs) and Pol II^1^. The evolutionary conserved TFIID complex plays a major role in transcription initiation as it is the first GTF to initiate the assembly of the pre-initiation complex (PIC) by recognizing the core promoter^2^. TFIID is a large multi-protein complex composed of the TATA box-binding protein (TBP) and 13 TBP-associated factors (TAFs) in metazoa^3^. The model suggesting that transcription is always regulated by the same transcription complexes has been challenged in metazoans by the discovery of cell-type specific complexes containing specialized GTF-, TBP- or TAF-paralogs^4^. Two TBP paralogues have been described in vertebrates: TBPL1 (TBP like factor; TLF, also known as TRF2) has been identified in all metazoan species^5-10^, while TBPL2 (also known as TRF3 or TBP2) has only been described in vertebrates^11,12^. Remarkably, while *Tbpl1* and *Tbpl2* mutants display embryonic phenotypes in non-mammalian species^7–10,12,13^, *Tbpl1* and *Tbpl2* loss of function in mouse results in male and female sterility, respectively^14-16^, suggesting that in mammals, these two TBP-like proteins are involved in cell specific transcription. While TBPL2 shares high degree of identity (92%) within the conserved saddle-shaped C-terminal DNA binding core domain of TBP^17^, the C-terminus of TBPL1 is more distant with only 42% identity^12^. A consequence is that TBPL2, but not TBPL1, is able to bind canonical TATA-box sequences *in vitro*^5,12,18^. The N-terminal domains of the three vertebrate TBP related factors do not show any conservation. All three vertebrate TBP related factors can interact with the GTFs TFIIA and TFIIB, and can mediate Pol II transcription initiation *in vitro*^12,13,18–20^. However, how alternative initiation complexes form, how they regulate cell type-specific transcription and how they recognize promoter sequences remain unknown.

Mapping of transcription start sites (TSSs), at single-nucleotide by Cap Analysis of Gene Expression (CAGE) revealed two main modes for transcription start site (TSS) usage^21^. Transcription initiation within a narrow region, called “sharp” (or focused) TSS-type, is common in highly active, tissue-specific gene promoters containing TATA boxes, while transcription initiation with multiple initiation positions within an about 100 bp region, called “broad” TSS promoter architecture^21^, are more characteristic to ubiquitously expressed and developmental regulated genes (reviewed in ^22^). During zebrafish maternal to zygotic transition it was described that two TSS-defining grammars coexist, in core promoters of constitutively expressed genes to enable their expression in the two regulatory environments^23^. Maternally active promoters in zebrafish tend to be sharp, with TATA-like, AT-rich (W-box) upstream elements guiding TSS selection, while embryonically active broad promoter architectures of the same genes appear to be regulated by nucleosome positioning ques. Although a number of germ cell specific as well as somatic transcriptional regulators have been well characterized during folliculogenesis (reviewed in ^24^), the exact actors and mechanisms required for setting up the oocyte specific transcriptome have not yet been identified in vertebrates.

Female germ cells develop during oogenesis leading to the formation of a highly differentiated and specialised cell, the oocyte. In females, oocytes enter meiosis during embryonic life. Quiescent primordial follicles composed of meiotically arrested oocytes at the late diplotene stage and surrounded by granulosa cells, are formed perinatally in mice (reviewed in ^24^). Shortly after birth, some primordial follicles enter folliculogenesis and undertake a growth phase during which a specific oocyte-specific transcriptome is set up, oocytes increase their size until the pre-antral follicular stage^25^. A remarkable feature of oocyte is the very high expression of retrotransposons driven by Pol II transcription. These elements are interspersed repetitive elements that can be mobile in the genome. One of the 3 major classes of retrotransposons in mammals is the long terminal repeat (LTR) retrotransposons derived from retroviruses, also known as endogenous retroviruses (ERVs) that is subdivided in 3 sub-classes: ERV1, ERVK and endogenous retrovirus like ERVL-MaLR (mammalian apparent LTR retrotransposons) (reviewed in ^26^). Transcription of mobile elements in specific cell types depends on the presence of a competent promoter recognition transcription machinery and/or the epigenetic status of the loci where these elements have been incorporated. Remarkably, MaLRs encode no known proteins, but MaLR-dependent transcription is key in initiating synchronous developmentally regulated transcription to reprogram the oocyte genome during growth^27^.

Remarkably, during oocyte growth TBP is absent and replaced by TBPL2^28^. Indeed, TBP is only expressed up to the primordial follicular oocytes and becomes undetectable at all subsequent stages during oocyte growth. In contrast, TBPL2 is highly expressed in the growing oocytes, suggesting that TBPL2 is replacing TBP for its transcription initiating functions during folliculogenesis^28^. In agreement with its oocyte-specific expression, a crucial role of TBPL2 for oogenesis was demonstrated in *Tbpl2*^*−/−*^ females, which show sterility due to defect in secondary follicle production^16,29^. In the absence of TBPL2, immunofluorescent staining experiments showed that elongating Pol II and histone H3K4me3 methylation signals were abolished between the primary and secondary follicle stage oocytes suggesting that Pol II transcription was impaired^16^. Initially TBPL2/TRF3 was suggested to be expressed during muscle differentiation^30^, but this observation was later invalidated^16,29^. Altogether, the available data suggested that TBPL2 is playing a specialized role during mouse oocyte development. However, how does TBPL2 regulate oocyte-specific transcription and what is the composition of associated transcription machinery, remained unknown.

Here we demonstrate that in oocytes TBPL2 does not assemble into a canonical TFIID complex, while it stably associates with TFIIA. The observation that the oocyte specific deletion of *Taf7*, a TFIID specific TAF, does not influence oocyte growth and maturation, corroborates the lack of TFIID in growing oocytes. Our transcriptomics analyses in wild type and *Tbpl2*^*−/−*^ oocytes show that TBPL2 mediates transcription of oocyte-expressed genes, including mRNA destabilisation factor genes, as well as MaLR ERVs. Our transcription start site (TSS) mapping from wild-type and *Tbpl2*^*−/−*^ growing oocytes demonstrates that TBPL2 has a strong preference for TATA-like motif in gene core promoters driving specific sharp TSS selection. This is in marked contrast with TBP/TFIID-driven TATA-less gene promoters in preceding stages that have broad TSS architecture. Our results show a role for the TBPL2-TFIIA transcription machinery in a major transition of the oocyte transcriptome mirroring the maternal to zygotic transition that occurs after fertilization, completing a full germ line cycle.

## Results

### Oocyte-specific TBPL2/TFIIA complex distinct from TFIIDs

To characterize TBPL2-containing transcription complexes we prepared whole cell extracts (WCE) from growing 14 days post-natal (P14) mouse ovaries and analysed TBPL2-associated proteins by anti-mTBPL2 immunoprecipitation (IP) coupled to label free mass spectrometry (Fig. 1a, Supplementary Fig. 1a, b). To determine the stoichiometry of the composition of the immunoprecipitated complexes, normalized spectral abundance factor (NSAF) values were calculated^31^. In the anti-TBPL2 IPs we identified TFIIA-αβ and TFIIA-γ subunits as unique GTF subunits associated with TBPL2 (Fig. 1a, Supplementary Data 1). As ovaries contain many other non-oocyte cell types which express TBP, in parallel from the same extracts we carried out an anti-TBP IP. The mass spectrometry of the anti-TBP IP indicated that TBP assembles into the canonical TFIID complex in non-oocyte cells (Fig. 1b, Supplementary Data 2). As growing oocytes represent only a tiny minority of ovary cells, we further tested the TBPL2-TFIIA interaction by a triple IP strategy (Fig. 1c): first, we depleted TAF7-containing TFIID complexes with an anti-TAF7 IP, second, the remaining TFIID and SAGA complexes, which contain also shared TAFs^32^, were depleted with an anti-TAF10 IP using the anti-TAF7 IP flow-through as an input, third we performed an anti-TBPL2 IP on the anti-TAF7/anti-TAF10 flow-through fraction (Fig. 1d-f, Supplementary Data 3). The analysis of this third consecutive IP further demonstrated that TBPL2 forms a unique complex with TFIIA-αβ, and TFIIFA-γ, but without any TFIID subunits.

**Fig. 1.**
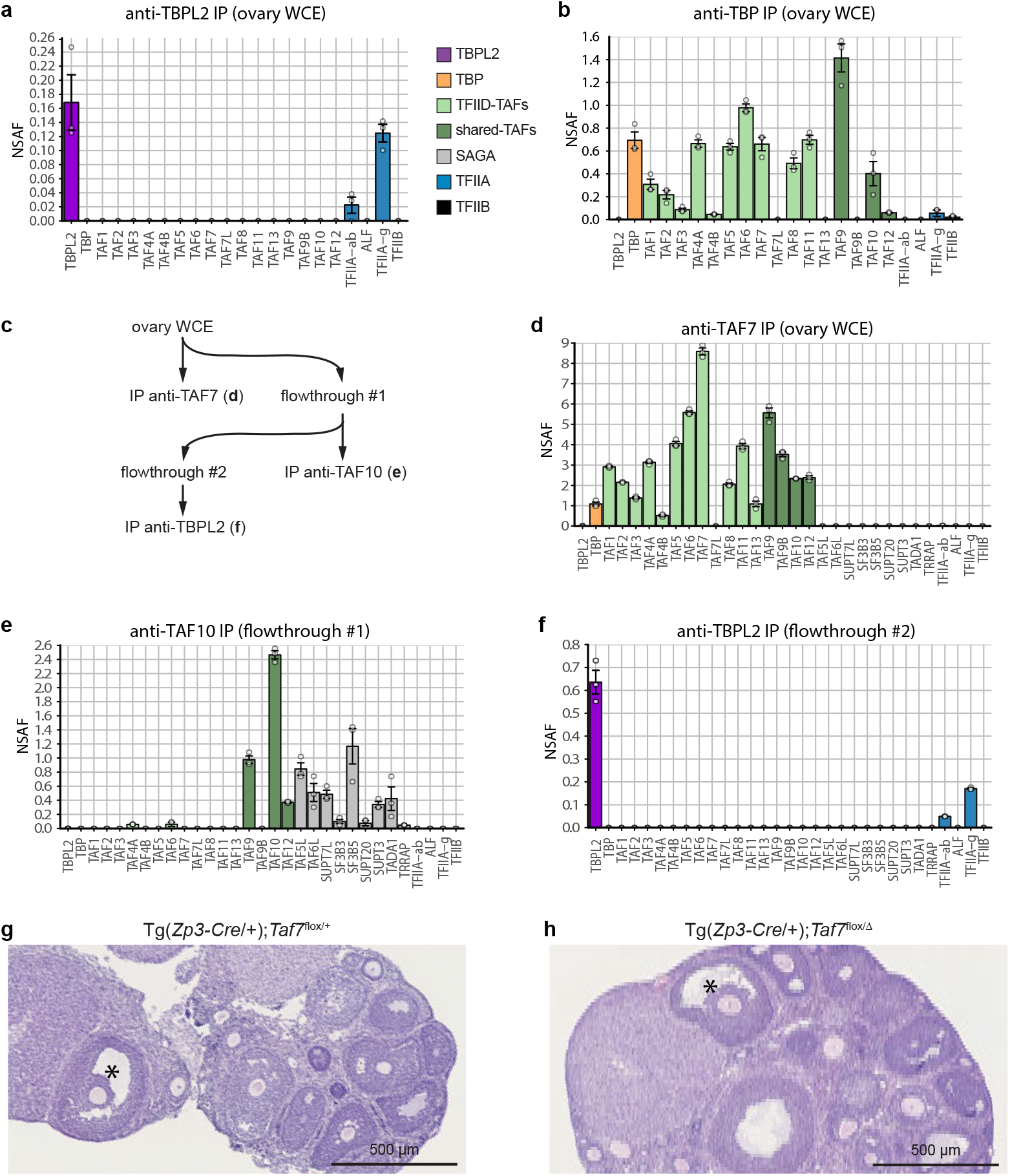
TBPL2 does not assemble in a TFIID-like complex in growing oocytes. **a** Anti-TBPL2 immunoprecipitation followed by mass spectrometry (IP-MS) analysis from three biological replicates of mouse ovarian whole cell extracts (WCE). The color code for the different proteins or complexes is indicated on the right. NSAF; normalized spectral abundance factor. **b** Anti-TBP IP-MS from ovarian WCE (three technical triplicates). The color code is the same as in (**a**). **c-f** Sequential IP-MS experiment from ovarian WCE (three technical triplicates). Strategy of the sequential immunoprecipitation (**c**), anti-TAF7 IP-MS (**d**), followed by an anti-TAF10 IP-MS (**e**) and then an anti-TBPL2 IP-MS (**f**). The color code is the same as in (**a**). **g, h**Representative views of haematoxylin and eosin stained ovaries section from control (**e**) and oocyte-specific *Taf7* mutant (Tg(*Zp3-Cre*/+);*Taf7*^flox/Δ^, **f**) ovaries (analyses of 3 sections from 3 biological replicates). The presence of antral follicles is indicated by an asterisk. Scale bars: 500 μm. Grey dots indicate replicates and error bars, +/− standard error of the mean in (**a**, **b**, **d-f**).

To further analyse the requirement of TFIID during oocyte growth, we carried out a conditional depletion of TFIID-specific *Taf7* gene during oocyte growth using the *Zp3-Cre* transgenic line^33^ (Supplementary Fig. 1c-g). Remarkably, TAF7 is only detected in the cytoplasm of growing oocyte (Supplementary Fig. 1c). The oocyte-specific deletion of *Taf7* did not affect the presence of secondary and antral follicles and the numbers of collected mature oocytes after superovulation (Fig. 1g, h, Supplementary Fig. 1f). The lack of phenotype is not due to an inefficient deletion of *Taf7*, as TAF7 immunolocalization is impaired (Supplementary Fig. 1d, e), and as oocyte-specific *Taf7* mutant females are severely hypofertile (Supplementary Fig. 1g). The observations that TBP is not expressed in growing oocytes, and that the oocyte specific deletion of *Taf7* abolishes the cytoplasmic localization of TAF7, but does not influence oocyte growth, show that canonical TFIID does not assemble in the nuclei of growing oocytes. Thus, our results together demonstrate that during oocyte growth a stable TBPL2-TFIIA complex forms, and may function differently from TBP/TFIID.

In order to further characterize the composition of the TBPL2-TFIIA complex, we took advantage of NIH3T3 cells artificially overexpressing TBPL2 (NIH3T3-II10 cells^28^). In this context where TBP and TAFs are present, TFIID is efficiently pulled down by an anti-TBP IP, but no interaction with TFIIA could be detected (Fig. 2a). Interestingly, the anti-TBPL2 IP showed that the artificially expressed TBPL2 can incorporate in TFIID-like complexes as TAFs were co-IP-ed (Fig. 2a), however with much lower stoichiometry (NSAF values) than that of TBP (Fig. 2b). In contrast, strong interaction with TFIIA-αβ and TFIIFA-γ were detected, suggesting that the TBPL2-TFIIA complex can be formed in the NIH3T3-II10 cells and that TBPL2, to the contrary to TBP has the intrinsic ability to interact with TFIIA. Remarkably, in spite of the high similarity between the core domains of TBP and TBPL2, no interaction with SL1 (TAF1A-D) and TFIIIB (BRF1) complexes that are involved in Pol I and Pol III transcription, respectively^34^) could be detected in the anti-TBPL2 IP, to the contrary to the anti-TBP IP in the NIH3T3-II10 (Fig. 2a, b) as well as in the ovary (Supplementary Data 1, 2), suggesting that TBPL2 is not involved in Pol I and Pol III transcription initiation in the growing oocytes.

**Fig. 2.**
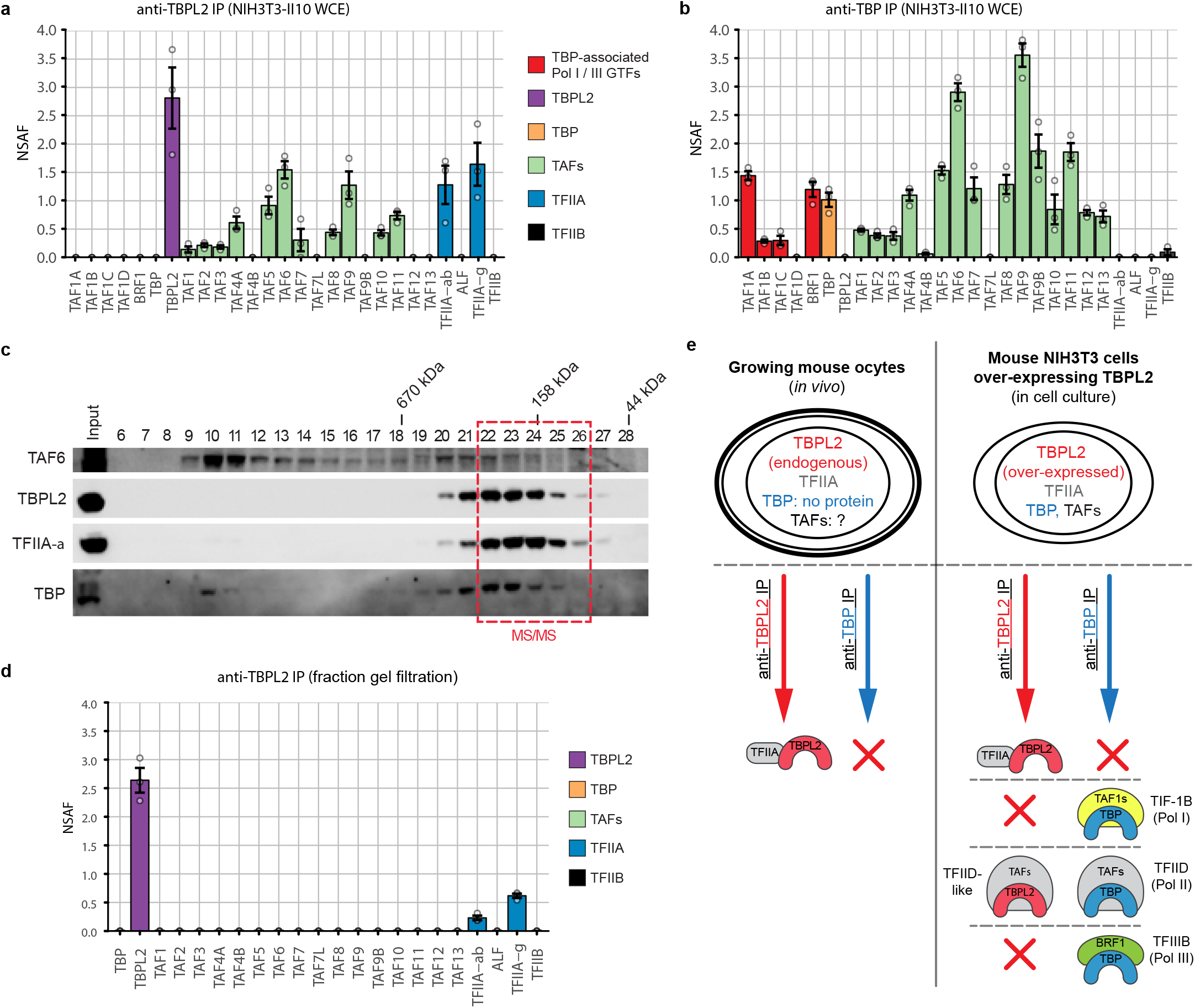
TBPL2 assembles into a TBPL2/TFIIA and TFIID-like complexes in NIH3T3 cells over-expressing TBPL2. **a** Anti-TBPL2 immunoprecipitation followed by mass spectrometry (IP-MS) analysis (three technical replicates) of NIH3T3 over-expressing TBPL2 (NIH3T3-II10) whole cell extracts (WCE). The color code for the different proteins or complexes is indicated on the right. **b** Anti-TBP IP-MS analysis (three technical replicates) of NIH3T3-II10 WCE. Color legend for the different proteins is the same as in (**a**). **c** Western blot of a Superose 6 gel filtration analysis of NIH3T3-II10 WCE probed with anti-TAF6 (top panel), anti-TBPL2 and anti-TFIIA-α (middle panels), and anti-TBP (bottom panel) antibodies. Fraction numbers are shown above each lane and the elution of known molecular mass markers is indicated above the panels. The pooled fractions used for mass spectrometry analysis are indicated in red. **d** Anti-TBPL2 IP-MS analysis (three technical replicates) of the gel filtration fraction indicated in (**c**). The color code for the different proteins or complexes is indicated on the right. NSAF; normalized spectral abundance factor. **e** Schematic representation of the fundamental differences existing between TBPL2- and TBP-containing complexes in growing oocytes and NIH3T3-II10 cells. Grey dots indicate replicates and error bars, +/− standard error of the mean in (**a**, **b**, **d**).

To analyze whether TBPL2 associates with TFIID TAFs and TFIIA in the same complex, we performed a gel filtration analysis of NIH3T3-II10 WCE. The profile indicated that most of the TBPL2 and TFIIA could be found in the same fractions (22-26) eluting around 150-200 kDa, while TBPL2 protein was below the detection threshold of the western blot assay in the TAF6-containing fractions 9-15 (Fig. 2c). To verify that TBPL2 and TFIIA are part of a same complex in fractions 22-26, we IP-ed TBPL2 from these pooled fractions and subjected them to mass spectrometric analysis. Our data confirmed that in these fractions eluting around 170 kDa, TBPL2 and TFIIA form a stable complex that does not contain any TAFs (Fig. 2d, supplementary Data 4). Thus, all these experiments together demonstrate that TBPL2/TFIIA form a stable complex in oocytes, where TBP is not expressed and TBPL2/TFIIA is the only promoter recognizing transcription complex that could direct Pol II transcription initiation (see the summary of all these IPs in Fig. 2e).

### TBPL2-dependent oocyte transcriptome

To characterize the growing oocyte-specific transcriptome and its dependence on TBPL2, we have performed a transcriptomic analysis of wild-type (WT) and *Tbpl2*^*−/−*^ oocytes isolated from primary (P7) and secondary follicles and (P14) (Fig. 3, 4, Supplementary Fig. 2, Supplementary Data 5). We observed down-regulation of a high number of oocyte-specific genes, out of which *Bmp15* and *Gdf9* served as internal controls^35,36^, as they were already described to be regulated by TBPL2^16^ (Fig. 3a, b and Supplementary Fig. 2a). Principal component analysis showed that the four distinct RNA samples clustered in individual groups and that the main explanation for the variance is the genotype, and then the stage (Supplementary Fig. 2b). Comparison of the RNA level fold changes between mutant and WT oocytes showed that in *Tbpl2*^*−/−*^, there is a massive down-regulation of the most highly expressed transcripts, both at P7 and P14 (Supplementary Fig. 2c). The Pearson correlation between the P7 and P14 fold change data sets for transcripts expressed above 100 normalized reads was close to 0.8 (Supplementary Fig. 2c), indicating that *Tbpl2* loss of function similarly altered RNA levels at P7 and P14 stages. We therefore focused on the P14 stage for the rest of the study.

**Fig. 3.**
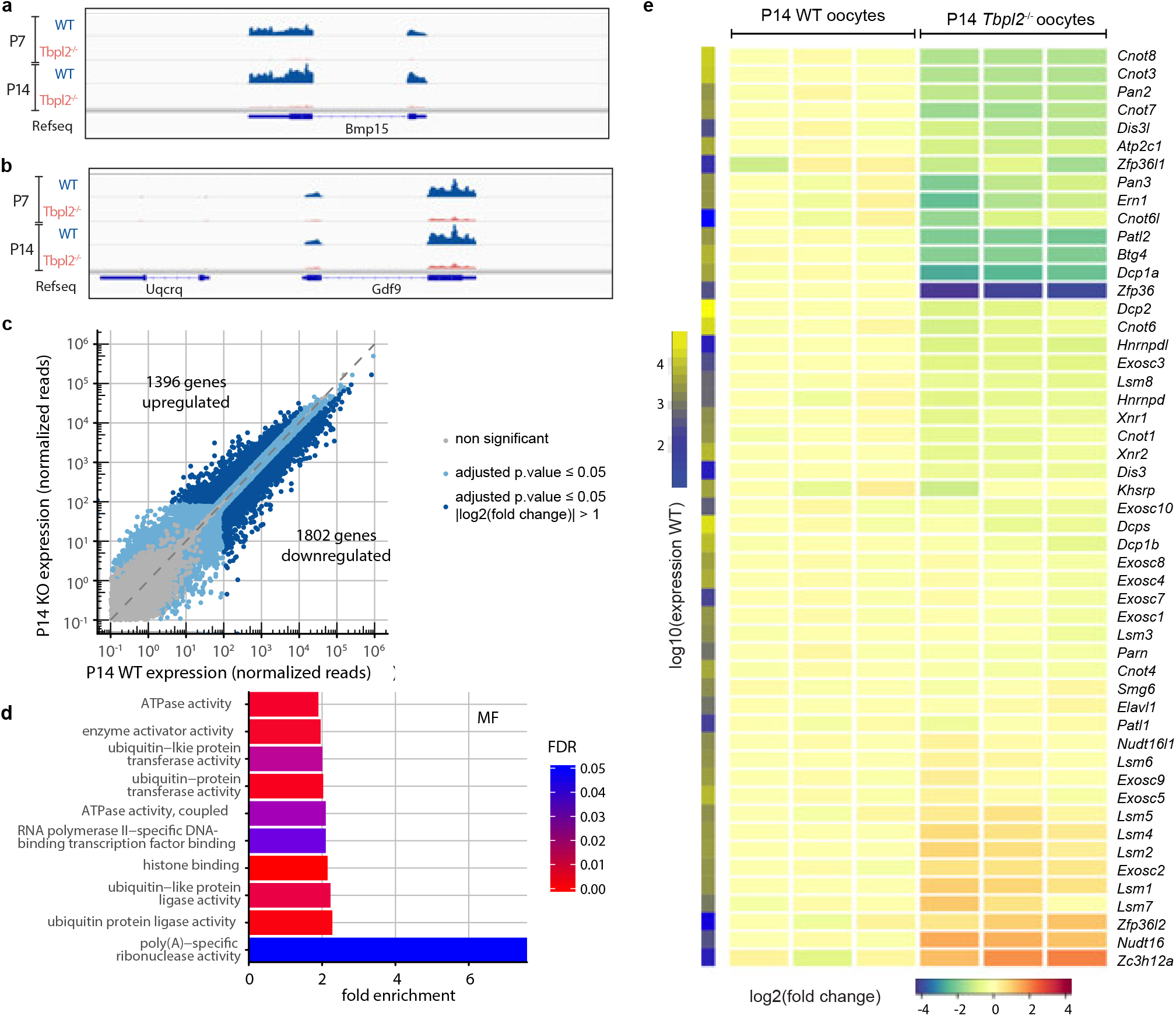
Expression of genes related to the mRNA deadenylation/decapping/decay pathways in growing *Tbpl2*^−/−^ mutant oocytes. **a, b** Normalized Integrative Genomic Viewer (IGV) snapshots of *Bmp15* (**a**) and *Gdf9* (**b**) loci. Exons and introns are indicated. **c** Expression comparison between wild-type (WT) and *Tbpl2*^−/−^ mutant post-natal day 14 (P14) oocytes (biological triplicates). Expression was normalized to the median size of the transcripts in kb. Grey dots correspond to non-significant genes and genes with high Cook’s distance, light-blue dots to significant genes for an adjusted p value ≤ 0.05 and dark-blue dots to significant genes for an adjusted p value ≤ 0.05 and an absolute log2 fold change > 1, after two-sided Wald test and Benjamini-Hochberg correction for multiple comparison. The number of up-or down-regulated genes is indicated on the graph. **d** Down-regulated genes GO category analyses for the molecular functions (MF). The top ten most enriched significant GO categories for a FDR ≤ 0.05 are represented. **e** Heatmap of selected genes involved in mRNA decay, decapping or deadenylation pathways. Expression levels in fold-change (compared to the mean of WT) of three biological replicates of P14 WT and P14 *Tbpl2*^−/−^ mutant oocytes are indicated. The fold change color legend is indicated at the bottom. The first column on the left corresponds to the log10 of expression, scale on the left.

**Fig. 4.**
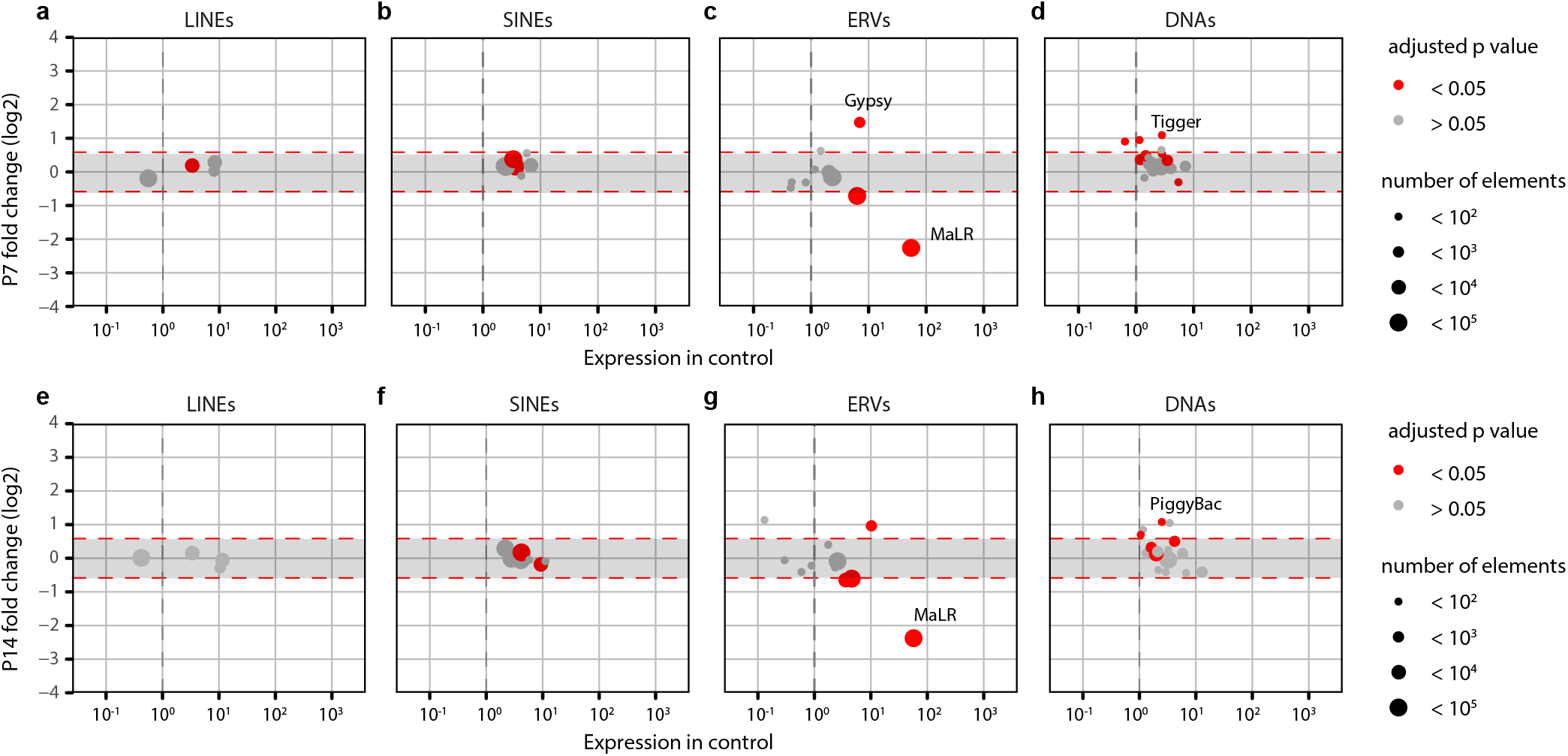
Expression of the mammalian apparent LTR retrotransposons (MaLR) endogenous retroviral elements (ERV) are down regulated in growing *Tbpl2*^−/−^ mutant oocytes. **a-h** Differential expression between wild type and *Tbpl2*^−/−^ mutant post-natal day (P) 7 (**a-d**) and P14 (**e-h**) oocytes of the different transposon classes; RNA transposon classes [LINEs (**a**, **e**), SINEs (**b**, **f**) and ERVs (**c**, **g**)] and DNA transposons (DNAs (**d**, **h**)). The ERV sub-class III mammalian apparent LTR retrotransposon (MaLR) family is the most severely affected in *Tbpl2*^−/−^ mutant oocytes at P7 and P14. Dot size and color are explained on the right side, adjusted *p* value after two-sided Wald test and Benjamini-Hochberg correction for multiple comparison.

In WT P14 oocytes transcripts corresponding to 10791 genes were detected. Importantly, many of these detected transcripts have been transcribed at earlier stages and are stored in growing oocytes^37^. As there is no Pol II transcription in *Tbpl2*^−/−^ growing oocytes^16^, RNAs detected in the *Tbpl2*^−/−^ mutant oocytes represent mRNAs transcribed by a TBP/TFIID-dependent mechanism and deposited into the growing oocytes independently of TBPL2 activity at earlier stages, i.e. at primordial follicular stage, where TBP is still expressed. The proportion of genes (1396) upregulated following *Tbpl2* deletion (Fig. 3c) can be explained by two mutually not exclusive ways: i) the consequence of the normalization to the library size resulting in a slight over-estimation of up-regulated transcripts, and ii) under-estimation of down-regulated transcripts and/or by transcript buffering mechanisms due to mRNA stabilization^38^. Validation of the up-regulation of some candidate transcripts levels (Supplementary Fig. 2d, e) strongly supports the latter hypothesis (but see also the next paragraph).

Nevertheless, we detected 1802 significantly downregulated transcripts in the *Tbpl2*^−/−^ oocytes (Fig. 3c). Key genes known to be expressed during oocyte growth, such as *Bmp15, Eloc, Fgf8, Gdf9* and *Zar1*^35,36,39^, were confirmed by RT-qPCR to be down-regulated (Supplementary Fig. 2f, g). These results suggest that TBPL2 has an important role in gene expression in the growing oocytes. Gene Ontology (GO) analyses of biological process of the identified down regulated categories of genes (Supplementary Data 6) indicated that many genes, involved in meiosis II and distinct cell cycle processes, were significantly down-regulated (Supplementary Fig. 2h). The most enriched molecular function GO category was “poly(A)-specific ribonuclease activity” containing many genes coding for factors or subunits of complexes contributing to deadenylation/decapping/decay activity in eukaryotes (Fig. 3d) (i.e. CCR4-NOT, PAN2/PAN3^40^; DCP1A/DCP2^41^, or BTG4^39^). In good agreement with the transcriptome analyses, transcripts coding for these “poly(A)-specific ribonuclease activity” factors were significantly down regulated in *Tbpl2*^−/−^ mutant P14 oocytes when tested by RT-qPCR (Fig. 3e, Supplementary Fig. 2i). Thus, in P14 oocytes TBPL2 is regulating the transcription of many genes coding for factors, which are in turn crucial in regulating the stability and translation of the mRNA stock deposited during early oogenesis, as well as transcription of meiosis II- and cell cycle-related genes to prepare the growing oocytes for the upcoming meiotic cell division.

A remarkable feature of oocyte is the very high expression of retrotransposons driven by Pol II transcription (see Introduction). As expected, in WT P7 and P14 oocytes the expression of ERVs was found to be the most abundant^27,42^ (Supplementary Fig. 3a-c). Importantly, the transcription of the vast majority of MaLR elements was the most affected in *Tbpl2*^*−/−*^ mutant oocytes at P7 and P14 (Fig. 4). Among them, three highly expressed members, *MT-int*, *MTA_Mm*, and *MTA_Mm-int*, were dramatically down-regulated in P7 and P14 *Tbpl2*^*−/−*^ mutant oocytes (Supplementary Fig. 3d, e). As in P14 oocytes TBPL2 depletion is reducing transcription more than 4-fold from MaLR ERVs, which often serve as promoters for neighbouring genes^27,42^, TBPL2 could seriously deregulate oocyte-specific transcription and consequent genome activation.

Therefore, this is the first demonstration that TBPL2 is orchestrating the *de novo* restructuration of the maternal transcriptome and that TBPL2 is crucial for indirectly silencing the translation of the earlier deposited TBP-dependent transcripts.

### TBPL2-driven promoters contain TATA box and are sharp

The promoter usage changes during zebrafish maternal to zygotic transition revealed different rules of transcriptional initiation in oocyte and in embryo, driven by independent and often overlapping sets of promoter “codes” ^23^. Importantly, this switch has not yet been demonstrated in mammals and the role of TBPL2 in this switch during oogenesis remained to be investigated. To this end, we mapped the TSS usage by carrying out super-low input carrier-CAGE (SLIC-CAGE) ^43^ from WT and *Tbpl2*^−/−^ P14 oocytes. To characterize only the TBPL2-driven promoters, we removed the CAGE tags present in the *Tbpl2*^−/−^ dataset from the WT P14 dataset, to eliminate transcripts that have been deposited at earlier stages (hereafter called “TBPL2-dependent”). Conversely, the *Tbpl2*^−/−^ dataset corresponds to the TBP/TFIID-dependent, or TBPL2-independent TSSs (hereafter called “TBPL2-independent”).

Next, we analysed the genome-wide enrichment of T- and/or A-rich (WW) dinucleotide motifs within the −250/+250 region centred on the dominant TSSs of the TBPL2-specific-only and TBPL2-independent-only oocyte TSS clusters (Fig. 5a, b). TBPL2-dependent TSS clusters are strongly enriched in a well-defined WW motif around their −30 bp region (Fig. 5a, red arrowhead) ^44^. In contrast, only about 1/3^rd^ of the TBPL2-independent TSS clusters contained WW-enriched motifs at a similar position (Fig. 5b, red arrowhead), as would be expected from promoters that lack maternal promoter code determinants^23,44^. As canonical TATA boxes are often associated with tissue-specific gene promoters, we investigated whether the above observed WW motif densities correspond to TATA boxes using the TBP position weight matrix (PWM) from the JASPAR database as a reference. To this end the presence of TATA boxes was analysed in the TSS clusters of the two data sets and revealed that TBPL2-dependent TSS clusters were enriched in high quality TATA boxes, including a clear increase in the proportion of canonical TATA boxes, when compared to TBPL2-independent TSS clusters (Fig. 5c). Genome browser view snapshots indicate that TSS clusters in P14 WT oocytes tend to be sharp and are associated with TATA-like motifs (Supplementary Fig. 4a, b). Analysis of the global distribution of the number of TSSs and of the width of the TSS clusters in the above defined two categories confirmed that TBPL2-dependent TSS are sharper compared to the TBPL2-independent TSS clusters (Supplementary Fig. 4c, d).

**Fig. 5.**
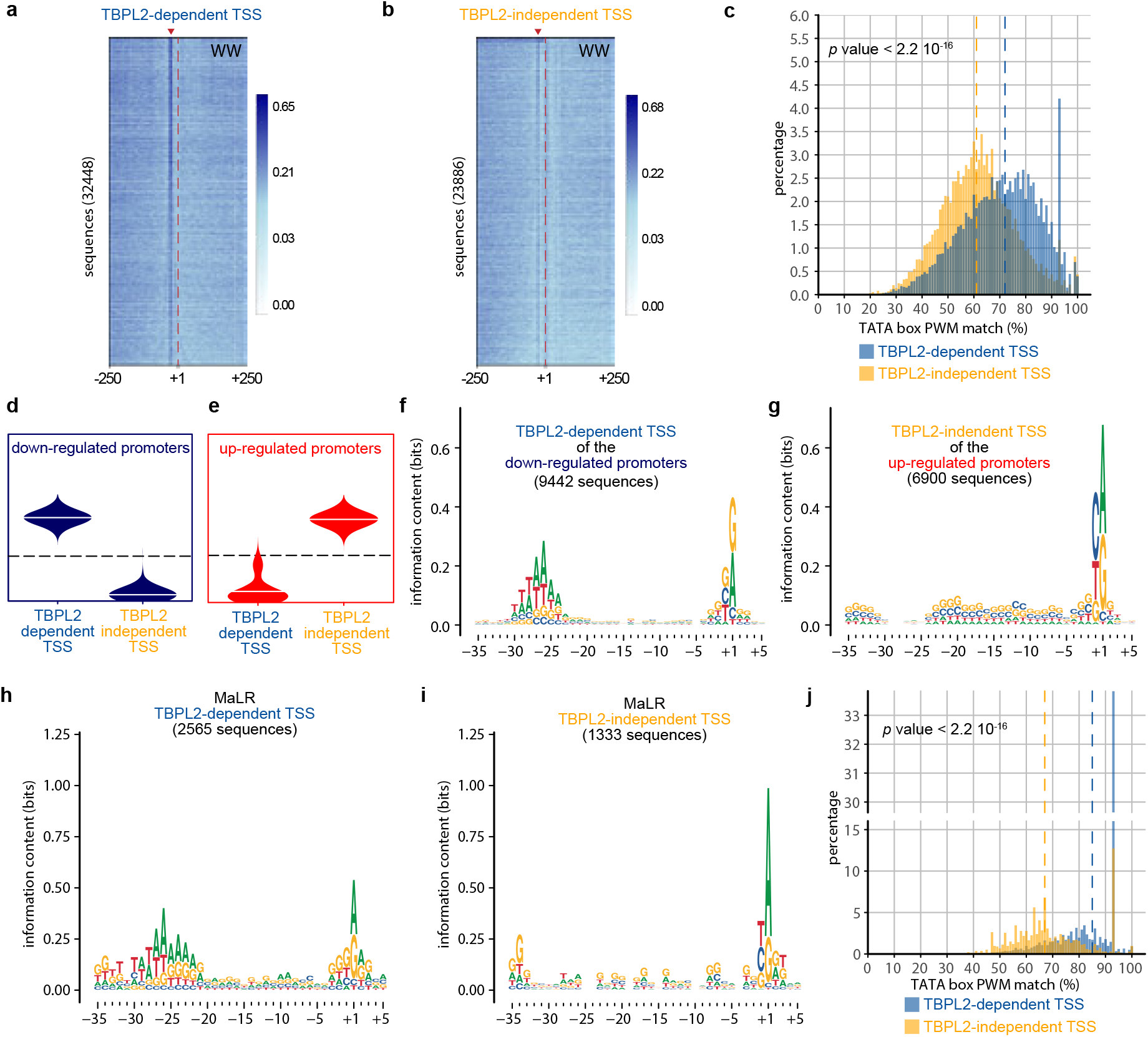
Core promoter regions of TBPL2 specific transcription units in post-natal day 14 oocytes are enriched in TATA-like elements and are sharp. **a, b** Genome-wide A/T-rich dinucleotide (WW) motif analyses of −250/+250 sequences centred on the dominant transcription start sites (TSS, position +1, dashed red line) of TBPL2-dependent (**a**, n = 32448) and TBPL2-indendent (**b**, n = 23886) TSS clusters. Sequences have been ordered by increasing size of the interquantile width of each. The red arrowheads indicate the WW enrichment at position −30 in the TBPL2-dependent TSS clusters (**a**) and the equivalent position in the TBPL2-independent TSS clusters (**b**). The number of TSS clusters is indicated in brackets. **c** Distribution of the best TATA box position weight matrix (PWM) matches within a −35 to −20 region upstream of the dominant TSSs (+1) of TBPL2-dependent (blue) compared to the TBPL2-independent (orange) TSS clusters. The dashed lines indicate the median of the TATA box PWM matches for the TBPL2-dependent (blue) and the TBPL2-independent (orange) TSS clusters (*p* value after a two-tailed Wilcoxon rank-sum test). **d, e** Two selected self-organizing map (SOM) groups of the consensus TSS clusters: the down-regulated promoters (blue, **d**) and the up-regulated promoters (red, **e**) groups. **f, g** Sequence logos of the −35/+5 sequence of the TBPL2-dependent dominant TSSs from the down-regulated promoters (**f**) and of the TBPL2-independent dominant TSSs from the up-regulated promoters (**g**). **h** Distribution of the best TATA box PWM matches within a −35 to −20 region upstream of the TBPL2-dependent (blue) and TBPL2-independent (orange) mammalian apparent LTR retrotransposons (MaLR) endogenous retroviral elements (ERV) dominant TSS. The dashed lines indicate the median of the TATA box PWM matches for the TBPL2-dependent (blue) and the TBPL2-independent (orange) TSS clusters (*p* value after a two-tailed Wilcoxon rank-sum test). **i, j**Sequence logo of the −35/+5 sequence of the MaLR ERVs TBPL2-dependent (**i**) and TBPL2-independent (**j**) dominant TSSs.

In order to test whether TBPL2 controls transcription initiation from maternal promoter code determinants, we grouped the expression profiles corresponding to each consensus TSS clusters, to characterise promoter activity profiles among datasets by performing self-organizing maps (SOM) ^45^ (Supplementary Fig. 4e). We then focussed on the two most distinct SOM groups: the down-regulated promoters (blue group, containing 9442 consensus TSS clusters) (Fig. 5d) and the up-regulated promoters (red group, with 6900 consensus TSS clusters) (Fig. 5e). Motif analyses of these two categories of promoters in their −35/+5 regions relative to the different dominant TSSs indicated that only the core promoters associated to TBPL2-dependent dominant TSSs belonging to the down-regulated gene promoters contain a well-defined 7 bp long TATA box-like motif (W-box) in their −31 to −24 regions (Fig. 5f, g, Supplementary Fig. 4f-i). Importantly, W-box associated TSSs architecture usage distribution for these TBPL2-dependent dominant TSSs was sharp (Supplementary Fig. 4j, l), as expected for motif-dependent transcriptional initiation^23,44^. In contrast, TBPL2-independent TSSs belonging to the up-regulated promoters exert a much broader TSS pattern (Supplementary Fig. 4k, m). Interestingly, GO analyses of the genes associated with the down-regulated promoters revealed a strong association with deadenylation/decapping/decay activity (Supplementary Fig. 4n-p, Supplementary Data 7), further confirming our initial RNA-seq analysis observations (Fig. 3).

Importantly, TSS architecture analyses of the TBPL2-dependent MaLR ERV TSSs indicated that the majority of MaLR core promoters contain high quality TATA box motif (median of the TATA box PWM match is 85%, Fig. 5h-j). These observations together demonstrate that the TBPL2/TFIIA complex drives transcription initiation primarily from core promoters that contain a TATA box-like motif in their core promoter and directs sharp transcription initiation from the corresponding promoter regions to overhaul the growing oocyte transcriptome.

In addition, we observed that TSS usage can shift within the promoter of individual genes depending on the genetic background (Supplementary Fig. 4b). To get more insights in these promoter architecture differences, we identified genome-wide 6429 shifting promoters by comparing either TBPL2-dependent to TBPL2-independent TSS data. These results are consistent with TSS shifts between somatic and maternal promoter codes occurring either in 5’, or 3’ directions (Fig. 6a, Supplementary Fig. 4q) ^44^. WW motif analysis indicated that on each shifting promoter, TBPL2-dependent dominant TSSs are associated with WW motifs, while TBPL2-independent dominant TSSs are not (Fig. 6b). In addition, the TATA box PWM match analyses indicated that these WW motifs are enriched in TATA box like elements compared to the corresponding TBPL2-independent shifting TSSs (Fig. 6c). Thus, our experiments provide a direct demonstration that TBP/TFIID and TBPL2/TFIIA machineries recognize two distinct sequences co-existing in promoters of the same genes with TBPL2 directing a stronger WW/TATA box-dependent sharp TSS selection in them.

**Fig. 6.**
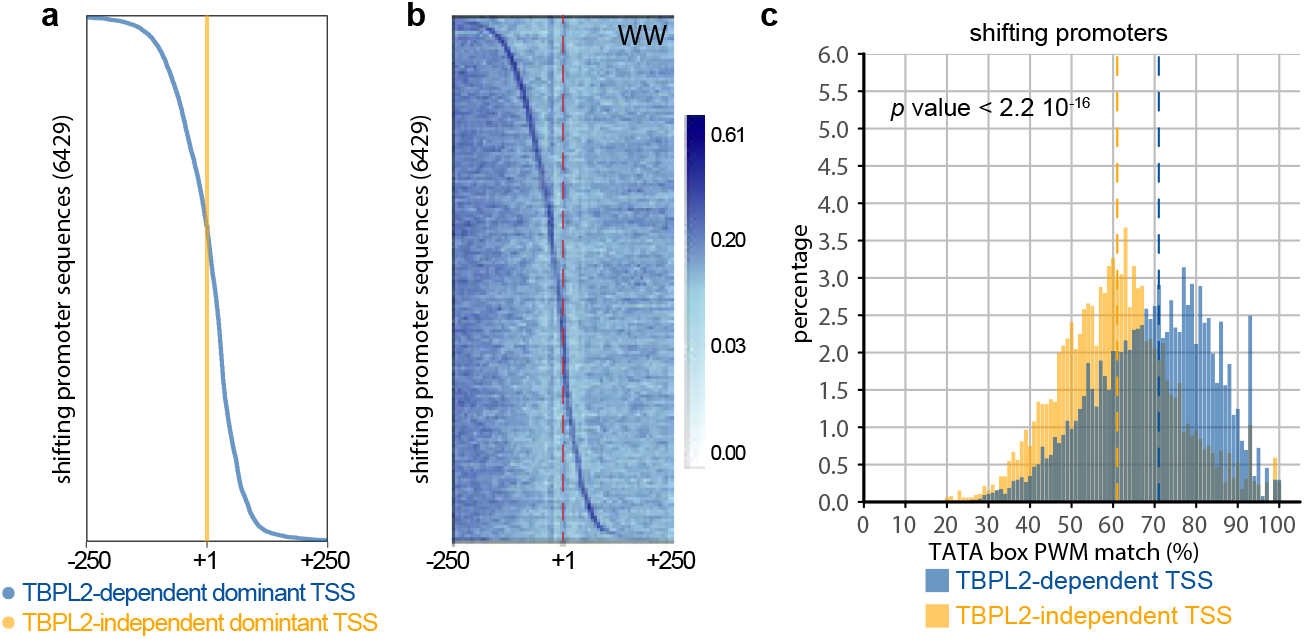
Shifting promoters. **a, b** Analysis of the TBPL2-dependent versus TBPL2-independent shifting promoters within a −250/+250 region centred on the position of the TBPL2-independent transcription start sites (TSS) clusters (position +1 in (**a, b**) and red dashed line in (**b**)). Position of the TBPL2-dependent (blue) and of the TBPL2-independent (orange) dominant TSSs for each shifting promoter sequence (**a**) and WW dinucleotide enrichment heat map (**l**) from the same set of sequences ordered as in (**b**). **c** Distribution of the best TATA box PWM matches within a −35 to −20 region upstream of the TBPL2-dependent (blue) and TBPL2-independent (orange) dominant TSS of the shifting promoters. The dashed lines indicate the median of the TATA box PWM matches for the TBPL2-dependent (blue) and the TBPL2-independent (orange) shifting TSS clusters (*p* value after a two-tailed Wilcoxon rank-sum test).

## Discussion

In this study, we showed that a unique basal transcription machinery composed of TBPL2 associated with TFIIA is controlling transcription initiation during oocyte growth, orchestrating a transcriptome change prior to fertilization using an oocyte-specific TTS usage.

TBPL2 expression in mouse is limited to the oocytes and in its absence, oocytes fail to grow and *Tbpl2*^−/−^ mouse females are sterile^16,28^. In a mirroring situation, TBPL1 (TRF2) expression is enriched during spermatogenesis and male germ cells lacking TBPL1 are blocked between the transition from late round spermatids to early elongating spermatids^14,15^. An interesting parallel between TBPL2 and TBPL1 is that both TBP-type factors form endogenous stable complexes with TFIIA. The beginning of TBPL2 accumulation in the oocyte nuclei, or TRF2 accumulation in male germ cell nuclei coincides with the phase of meiosis I^15,28,46^. It is thus conceivable that TBPL2-TFIIA in oocytes, or TRF2-TFIIA during spermatogenesis are involved in the control of gene expression in a meiotic context to set up the corresponding transcriptome. Interestingly, both transcription complexes seem to function in a compacted chromatin environment in which TBP/TFIID probably cannot. However, while TBPL2 and TBP show contrasting expression patterns in the oocytes^28^, TBPL1 and TBP are co-expressed in spermatids^46,47^ and it has been suggested that TBPL1 is a testis-specific subunit of TFIIA that is recruited to PIC containing TFIID and might not primarily act independently of TFIID/TBP to control gene expression in round spermatids^48^. While TBPL1 forms a complex also with the TFIIA-αβ paralogue, ALF, in testis^48–50^, TBPL2 does not stably associate with ALF, in spite of the fact that it is expressed in oocytes^50^.

TBP-like factors are bipartite proteins with variable N-terminal domains and a relatively well conserved shared C-terminal domains (core domains) forming a saddle-like structure with a concave surface that is known to bind to DNA^17^. Interestingly, TBPL1 has a very short N-terminal domain^5,18^ suggesting that it lost some abilities to interact with partners. Our data suggest that despite their very high similarity (92% identity between the core domains of TBP and TBPL2 (reviewed in ^51^), TBP and TBPL2 display different properties as they seem to recognize different DNA sequences to regulate gene promoters with different promoter architectures. Our IP-MS analyses from ovary WCE indicate that contrary to TBP, TBPL2 does not interact with TAFs in growing oocytes. Our analyses in the NIH3T3-II10 cells that overexpresses TBPL2 showed that TBPL2 can interact with TAFs in this artificial situation, albeit with less affinity compared to TFIIA, or TBP-TAFs interactions. Our transcriptomic data indicates that all *Taf* mRNAs, except *Taf7l* are detected in growing oocytes (Supplementary Data 5). However, whether they are also expressed in oocytes at the protein level is not yet known, except for TAF4B that has been detected in female neonate oocytes^52^. Nevertheless, our data suggest that TAF7 is expressed, but localized to the cytoplasm. It could be conceivable that, similarly to *Tbp* mRNA that is transcribed, but not translated in oocytes^53^, *Taf* mRNA translations (other than *Taf7*) are also inhibited and as a result the canonical TFIID, or its building blocks, cannot be assembled, and as a result the canonical TFIID is not present in the nuclei of growing oocytes. Another reason why TBPL2 does not interact with TAFs, or ALF, but rather interacts with TFIIA could be its N-terminal domain that is very different from TBP (only 23% identity^51^).

TBPL2 proteins from different vertebrates show a high degree of similarity in their C-terminal core domains amongst themselves; but display very little conservation in their N-terminal domains^12^. It is interesting to note that TBPL2 deficiency leads to an embryonic phenotypes in Xenopus^13^ and zebrafish^12^, because, contrary to the mouse, TBPL2 is still present in the embryo after fertilization and thus may act in parallel with TBP in the transcription of specific embryonic genes^10,54^. The molecular mechanism by which TBPL2 controls the transcription of these specific sets of genes in frogs and in fish has not been studied. In contrary, TBPL2 in mammals are only expressed in growing oocytes and the only phenotype that can be observed in mammals is female sterility^16,29^.

LTR retrotransposons, also known as ERVs, constitute ~10% of the mouse genome (reviewed in ^55^). While their expression is generally suppressed by DNA methylation and/or repressive histone modifications, a subset of ERV subfamilies retain transcriptional activity in specific cell types ^56^. ERVs are especially active in germ cells and early embryos (reviewed in 26). Indeed, many genome-wide transcripts are initiated in LTRs, such as for example of MaLRs in mouse oocytes, which constitute ~5% of the genome^57^. Members of the MT subfamily of MaLRs are particularly active in oocytes and hundreds of MT LTRs have been co-opted as oocyte-specific gene promoters^27,58^. As LTR-initiated transcription units shape also the oocyte methylome, it will be important to analyse also how TBPL2 influences DNA methylation in oocytes.

Oocytes display remarkable post-transcriptional regulatory mechanisms that control mRNA stability and translation. During oogenesis, the oocyte genome is transcriptionally active, and the newly synthesized maternal mRNAs are either translated or stored in a dormant form (reviewed in ^37^). The newly synthesized transcripts receive a long poly(A) tail and subsequently undergo poly(A) shortening in the oocyte cytoplasm, preventing translation. Until resumption of meiosis, mRNAs with a short poly(A) tail are stored in the cytoplasm in a dormant form (for a review, see ^59^). Thus, poly(A) tail deadenylation, amongst other activities, coordinates post-transcriptional regulation of the oocyte mRNA pool. Interestingly, TBPL2 is regulating the activity of several deadenylation/decapping/decay complexes and in the absence of TBPL2, we observed apparent stabilization of a significant number of transcripts, suggesting that in wild type oocytes TBPL2 is indirectly inhibiting the translation of mRNAs, and/or inducing the degradation of the mRNAs, previously deposited by TFIID/TBP in the primordial follicular oocytes (Fig. 7). To put in place the growing oocyte specific maternal transcriptome TBPL2 is controlling the production of new mRNAs using a maternal specific TSS grammar, as most of these transcripts will remain in the oocyte after transcriptional quiescence. Remarkably, as TBPL2 does not interact with Pol I and Pol III transcription machineries in the growing oocytes, this strongly suggest that rRNA and tRNA are deposited very early during oogenesis in amounts sufficient for the initiation of development.

**Fig. 7.**
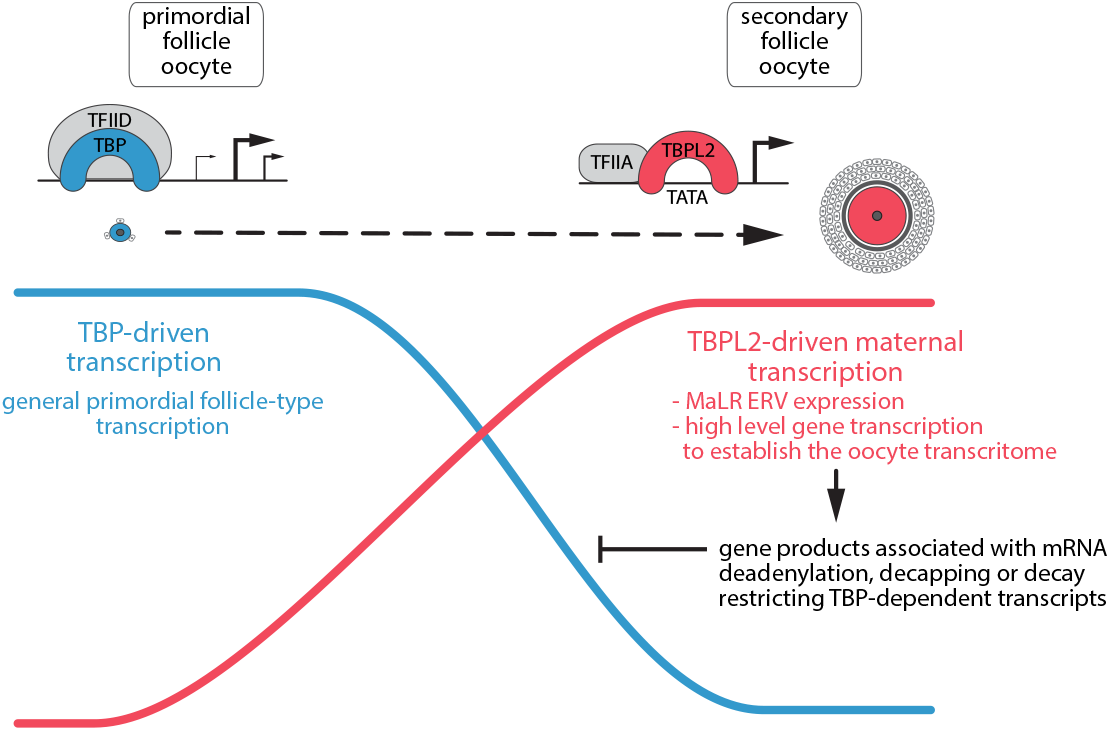
Transcriptome overhaul controlled by TBPL2/TFIIA during oocyte growth. At the beginning of oocyte growth, the transcriptome in primordial and early primary follicles (blue cell) depends on TFIID/TBP (blue complex) transcription from broad promoters (blue line). As TBP protein disappears, Pol II transcription initiation is mediated (red line) only by the oocyte-specific TFIIA/TBPL2 complex (red complex) from sharp promoters. At the growing oocyte stage (red cell) TFIIA/TBPL2 complex is responsible for initiating high levels of Pol II transcription, such as mammalian apparent LTR retrotransposons (MaLR) endogenous retroviral elements (ERV) expression and also expression of genes of which the gene products survey/limit the stability and/or translatability of transcripts previously deposited by TFIID/TBP-dependent Pol II transcription.

Therefore, it seems that TBPL2 contributes to establish a novel TBPL2-dependent growing oocyte transcriptome and consequent proteome required for further development and oocyte competence for fertilization (Fig. 7). The indirect regulation of previously deposited mRNAs by a global transcription regulator resembles the well characterised maternal to zygotic transition (MZT), during which a clearance of inherited transcriptome is mediated by *de novo* gene products generated by newly activated transcription machinery (reviewed in ^59^). At hundreds of gene promoters, two distinct TSS-defining “grammars” co-exist in close proximity genome-wide and are differentially utilised in either TBPL2/TFIIA in primary/secondary follicular oocytes, or by TBP/TFIID in primordial follicular oocytes or in the fertilized embryo. This again shows a striking parallel to MZT^23^, where multiple layers of information are embedded in the same promoter sequence, each representing a different type of regulatory grammar interpreted by dedicated transcription machinery depending on the cellular environment.

## Methods

### Cell lines and cell culture

The NIH3T3-II10 line overexpressing TBPL2 and the control NIH3T3-K2 have already been described^28^ and were maintained in high glucose DMEM supplemented with 10% of new-born calf serum at 37°C in 5% CO_2_.

### Whole cell extracts

NIH3T3-II10 and NIH3T3-K2 cells cultured in 15 cm dish were washed twice with 1x PBS, subsequently harvested by scrapping on ice. Harvested cells were centrifuged 1000 g at 4°C for 5 min and then resuspended in 1 packed cell volume of whole cell extraction buffer (20 mM Tris HCl pH7.5, 2 mM DTT, 20% Glycerol, 400 mM KCl, 1x Protease inhibitor cocktail (PIC, Roche)). Cell lysates were frozen in liquid nitrogen and thawed on ice for 3 times, followed by centrifugation at 20817 g, at 4°C for 15 min. The supernatant was collected and protein concentration was measured by Bradford protein assay (Bio-Rad). The cell extracts were used directly for immunoprecipitation and western blot, or stored at −80°C.

Ovaries collected from post-natal day 14 (P14) CD1 and C57BL/6N female mice were homogenized in whole cell extraction buffer [20 mM Tris HCl pH7.5, 2 mM DTT, 20% Glycerol, 400 mM KCl, 5x PIC (Roche)]. Cell lysates were frozen in liquid nitrogen and thawed on ice for 3 times, followed by centrifugation at 20817 g, at 4°C for 15 min. The supernatant extracts were used directly for immunoprecipitation.

### Antibodies and antibody purification

The antibodies are listed in Supplementary Table 1. The IGBMC antibody facility raised the anti-TBPL2 polyclonal 3024 serum against the CPDEHGSELNLNSNSSPDPQ peptide (amino acids 111-129) coupled to ovalbumin and injected into one 2 months-old female New-Zeland rabbit. The resulting serum was affinity purified by using the Sulfolink Coupling Gel (Pierce) following the manufacturer’s recommendations.

### Immunoprecipitation

Ovary extract were incubated with anti-GST (10 μg per IP), anti-TBP (10 μg per IP), anti-TBPL2 (3024, 12 μg (36 μg for gel filtration) per IP), anti-TAF7 (10 μg per IP) or anti-TAF10 (10 μg per IP) coated Dynabeads (Invitrogen) at 4°C overnight. After incubation, beads were washed 3 × 5min at 4°C with 500 mM KCl buffer [25 mM Tris-HCl (pH 7.9), 5 mM MgCl2, 10% glycerol, 0.1% NP40, 2 mM DTT, 500 mM KCl and 1x PIC (Roche)], then washed 3 × 5min at 4°C with 100 mM KCl buffer (25 mM Tris-HCl pH 7.9, 5 mM MgCl2, 10% glycerol, 0.1% NP40, 2 mM DTT, 100 mM KCl and 1x). Immunoprecipitated proteins were eluted with 0.1 M glycine pH 2.8 and neutralized with 1.5 M Tris-HCl pH 8.8.

Immunoprecipitation performed from whole cell extracts of NIH3T3-II10 and NIH3T3-K2 cells were following the same procedures with protein G Sepharose beads (GE Healthcare): 18 μg rabbit anti-TBPL2 (3024) and 15 μg anti-TBP per IP.

### Western blot analyses

Protein samples (15-25 μg of cell extracts or 15 μL of IP elution) were mixed with 1/4^th^ volume of loading buffer (100 mM Tris-HCl pH 6.8, 30% glycerol, 4% SDS, 0.2% bromophenol blue and freshly added 100 mM DTT) and boiled for 10 min. Samples were then resolved on a 10 % SDS-PAGE and transferred to nitrocellulose membrane (Protran, Amersham). Membranes were blocked in 3% non-fat milk in 1x PBS at room temperature (RT) for 30 min, and subsequently incubated with the primary antibody overnight at 4°C (dilution 1/1000). Membranes were washed three times (10 min each) with 1x PBS - 0.05% Tween20. Membranes were then incubated with HRP-coupled goat anti Mouse Ig (Jackson ImmunoResearch, #115-036-071, dilution 1/10000) or HRP-coupled goat anti-Rabbit Ig (Jackson ImmunoResearch, #115-035-144, dilution 1/10000) for 1 hour at RT, followed by ECL detection (Thermo Fisher). The signal was acquired with the ChemiDoc imaging system (Bio-Rad).

### Mass spectrometry analyzes and NSAF calculations

Samples were TCA precipitated, reduced, alkylated and digested with LysC and Trypsin at 37°C overnight. After C18 desalting, samples were analyzed using an Ultimate 3000 nano-RSLC (Thermo Scientific, San Jose, California) coupled in line with a linear trap Quadrupole (LTQ)-Orbitrap ELITE mass spectrometer via a nano-electrospray ionization source (Thermo Scientific). Peptide mixtures were loaded on a C18 Acclaim PepMap100 trap column (75 μm inner diameter × 2 cm, 3 μm, 100 Å; Thermo Fisher Scientific) for 3.5 min at 5 μL/min with 2% acetonitrile (ACN), 0.1% formic acid in H_2_O and then separated on a C18 Accucore nano-column (75 μm inner diameter × 50 cm, 2.6 μm, 150 Å; Thermo Fisher Scientific) with a 240 minutes linear gradient from 5% to 50% buffer B (A: 0.1% FA in H_2_O / B: 80% ACN, 0.08% FA in H_2_O) followed with 10 min at 99% B. The total duration was set to 280 minutes at a flow rate of 200 nL/min.

Proteins were identified by database searching using SequestHT with Proteome Discoverer 1.4 software (Thermo Fisher Scientific) a combined Mus musculus database generated using Uniprot [https://www.uniprot.org/uniprot/?query=proteome:UP000000589&sort=score] (Swissprot, release 2015_11, 16730 entries) where 5 interesting proteins sequences (TrEMBL entries: TAF4, ATXN7L2, TADA2B, BTAF1 and SUPT3) were added. Precursor and fragment mass tolerances were set at 7 ppm and 0.5 Da respectively, and up to 2 missed cleavages were allowed. Oxidation (M) was set as variable modification, and Carbamidomethylation© as fixed modification. Peptides were filtered with a false discovery rate (FDR) at 5 %, rank 1 and proteins were identified with 1 unique peptide. Normalized spectral abundance factor (NSAF)^31^ were calculated using custom R scripts (R software version 3.5.3). Only proteins detected in at least 2 out of 3 of the technical or biological replicates were considered for further analyses.

### Gel filtration

A Superose 6 (10/300) column was equilibrated with buffer consisting of 25mM Tris HCl pH 7.9, 5 mM MgCl2, 150 mM KCl, 5% Glycerol, 1 mM DTT and 1x PIC (Roche). Five hundred μL of whole cell extracts containing ∼5 mg of protein were injected in an ÄKTA avant chromatography system (Cytiva) and run at 0.4 mL/min. Protein detection was performed by absorbance at 280 nm and 260 nm. Five hundred μL fractions were collected and analysed by Western blot and IP-MS.

### Animal experimentation

Animal experimentations were carried out according to animal welfare regulations and guidelines of the French Ministry of Agriculture and procedures were approved by the French Ministry for Higher Education and Research ethical committee C2EA-17 (project n°2018031209153651). The Tg(*Zp3-Cre*), *Taf7*^flox^ and *Tbpl2*^−^ mouse lines have already been described^16,33,60^.

### Histology analyses of ovaries

Ovaries were collected from 6 weeks-old Tg(*Zp3-Cre*/+);*Taf7*^flox/+^ and Tg(*Zp3-Cre*/+);*Taf7*^flox/Δ^ oocyte specific mutant females, fixed in 4% paraformaldehyde (Electron Microscopy Sciences) over-night at 4°C, washed 3 times in PBS at room temperature and embedded in paraffin. Five μm-thick sections were stained with haematoxylin and eosin and images were acquired using a slide scanner Nanozoomer 2.0HT (Hamamatsu Photonics).

### Immunolocalization of TAF7 in the oocytes

Ovaries were dissected in PBS, fixed overnight in 4%PFA/PBS at 4°C, rinsed three times in PBS, equilibrated in 30% sucrose/PBS and embedded in Cryomatrix (Thermo Scientific) in liquid nitrogen vapor. Fifteen μm-thick sections were obtained on a Leica cryostat and stored at −80°C. Sections were rehydrated in TBS (50 mM Tris, 150 mM NaCl, pH7.5), and permeabilized with 0.5% Triton-X-100 (Sigma) and rinsed twice again in TBS before blocking in 3% BSA, 1% Goat serum, 0.1% Tween-20 (Sigma). Immunolabeling was then performed using M.O.M® Immunodetection Kit, Basic (Vector Laboratories, BMK-2202). Purified anti-TAF7 rabbit polyclonal antibody (dilution 1/300) was revealed using an Alexa Fluor 488 goat anti-rabbit IgG (Invitrogen #A-11108, dilution 1/1000). Sections were counterstained with DAPI (4′,6-diamidino-2-phenylindole dihydrochloride; Molecular Probes). Pictures were taken using a TCS SP5 Inverted confocal (Leica) with a 40x Plan APO objective (CX PL APO 40x/1.25-0.75 OIL CS) and analyzed using Fiji 2.0.

### Supervovulation

Five units of pregnant mare serum (PMS) was injected intraperitoneally in 4-week-old female mice between 2-4 pm. After 44-46 hours, GV oocytes were collected from the ovaries by puncturing with needles.

### Oocytes collection

After dissection, ovaries are freed from adhering tissues in 1x PBS. Series of 6 ovaries were digested in 500 μL of 2 mg/mL Collagenase (SIGMA), 0.025% Trypsin (SIGMA) and 0.5 mg/mL type IV-S hyaluronidase (SIGMA), on a ThermoMixer (Eppendorf) with gentle agitation for 20 minutes. The digestion was then stopped by the addition of 1 mL of 37°C pre-warmed αMEM - 5% FBS. The oocytes were then size-selected under a binocular.

### RNA preparation

Pool of 100-200 oocytes collected were washed through several M2 drops, and total RNA was isolated using NucleoSpin RNAXS kit (Macherey-Nagel) according to the user manual. RNA quality and quantity were evaluated using a Bioanalyzer. Between 5-10 ng of RNA was obtained from each pool of oocytes.

### RNA-seq analyses

PolyA+ RNA seq libraries were prepared using the SMART-Seq v4 UltraLow Input RNA kit (Clonetch) followed by the Nextera XT DNA library Prep kit (Illumina) according to the manufacturer recommendations from 3 biological replicates for each condition (P7 wild-type (WT), P7 *Tbpl2*^−/−^ mutant, P14 WT and P14 *Tbpl2*^−/−^ mutant oocytes) and sequenced 50 pb single end using an Illumina HiSeq 4000 (GenomEast platform, IGBMC).

Reads were preprocessed in order to remove adapter, polyA and low-quality sequences (Phred quality score below 20). After this preprocessing, reads shorter than 40 bases were discarded for further analysis. These preprocessing steps were performed using cutadapt version 1.10^61^. Reads were mapped to spike sequences using bowtie version 2.2.8^62^, and reads mapping to spike sequences were removed for further analysis. Reads were then mapped onto the mm10 assembly of *Mus musculus* genome using STAR version 2.7.0f^63^. Gene expression quantification was performed from uniquely aligned reads using htseq-count version 0.9.1^64^, with annotations from Ensembl version 96 and “union” mode. Read counts were normalized across samples with the median-of-ratios method, to make these counts comparable between samples and differential gene analysis were performed using the DESeq2 version 1.22.2^65^. All the figures were generated using R software version 3.5.3.

### RT-qPCR

Complementary DNA was prepared using random hexamer oligonucleotides and SuperScript IV Reverse Transcriptase (Invitrogen) and amplified using LightCycler® 480 SYBR Green I Master (Roche) on a LightCycler® 480 II (Roche). Primers used for qPCR analysis are listed in Supplementary Table 2.

### Repeat element analyses

Data were processed as already described^66^ using Bowtie1 (version 1.2.2) ^67^ instead of Maq. The repeatMasker annotation was used to identified the different types of repeat elements (Smit, AFA, Hubley, R & Green, P. RepeatMasker Open-4.0. 2013-2015 http://www.repeatmasker.org). Differential expression analyses were performed using DESeq2 (version 1.22.2) ^65^. All the figures were generated using R custom scripts (version 3.5.3).

### SLIC-CAGE analyses

Twenty-eight and 13 ng of total RNA isolated from P14 oocytes (biological replicate 1 and replicate 2, approximately 500-1000 oocytes pooled for each replicate) and 15 ng of total RNA isolated from P14 *Tbpl2*^−/−^ mutant oocytes (approximately 550 pooled oocytes) were used for SLIC-CAGE TSS mapping^43^. Briefly, 5 μg of the carrier RNA mix were added to each sample prior to reverse transcription, followed by the cap-trapping steps designed to isolate capped RNA polymerase II transcripts. The carrier was degraded from the final library prior to sequencing using homing endonucleases. The target library derived from the oocyte RNA polymerase II transcripts was PCR-amplified (15 cycles for P14 WT, 16 cycles for P14 *Tbpl2*^−/−^ mutant) and purified using AMPure beads (Beckman Coulter) to remove short PCR artifacts (< 200bp, size selection using 0.8 x AMPure beads to sample ratio). The libraries were sequenced using HiSeq2500 Illumina platform in single-end, 50 bp mode (Genomics Facility, MRC, LMS).

Sequenced SLIC-CAGE reads were mapped to the reference *M. musculus* genome (mm10 assembly) using the Bowtie2^62^ with parameters that allow zero mismatches per seed sequence (22 nucleotides). Uniquely mapped reads were kept for downstream analyses using CAGEr Bioconductor package (version 1.20.0) ^68^ and custom R/Bioconductor scripts. Bam files were imported into R using the CAGEr package, where the mismatching additional G, if added through the template-free activity of the reverse transcriptase, was removed. Same samples sequenced on different lanes and biological replicates were merged prior to final analyses.

### Promoter analyses

In order to consider only the CAGE TSS dependent only on TBPL2, we removed all the P14 WT CAGE tags at position where CAGE tags were also present in the P14 *Tbpl2*^*−/−*^ mutant CAGE tags dataset: for the rest of the analysis, this dataset was called “TBPL2-dependent” and we compared it to the P14 *Tbpl2*^*−/−*^ mutant CAGE data (hereafter called “TBPL2-independent”). Briefly, a CAGE set object was created from the TBPL2-dependent and TBPL2-independent CTSS files using CAGEr Bioconductor package (version 1.20.0) ^68^, data were normalized using normalizeTagCount (fitInRange = c(5,1000), alpha = 1.53, T = 1e6) and the powerLaw option. Cluster of CTSS were collected using clusterCTSS (threshold = 1, thresholdIsTpm = TRUE, nrPassThreshold = 1, method = “distclu”, maxDist = 20, removeSingletons = TRUE, keepSingletonsAbove = 5). Width of the TSS regions was calculated using cumulativeCTSSdistribution and quantilePositions (clusters = “tagClusters”, qLow = 0.1, qUp = 0.9): interquantile width corresponds to the 10^th^-90^th^ percentile of the total tag cluster signal. In order to compare the different samples, consensus promoters were computed using aggregateTagCluster (tpmThreshold = 3, qLow = 0.1, qUp = 0.9, maxDist = 100). Self-organizing map (SOM) expression profiling was performed using getExpressionProfiles using a tpmThrshold of 3, the method “som”, xDim = 3 and yDim = 2. Shifting TSS were obtained after calculation of the cumulative distribution along the consensus clusters using cumulativeCTSSdistribution and calculation of the shift score using scoreShift with the Kolomogorov-Smirnov test. Shifting promoters were extracted using getShiftingPromoters (tpmThreshold = 3, scoreThreshold = −Inf, fdrThreshold = 0.01). TSSs corresponding to the MaLR-ERVS were identified after annotation using HOMER (version 4.10) ^69^.

Sequences analyses were performed using Bioconductor R seqPattern (version 1.14) and R custom scripts. WW dinucleotides enrichment was computed using plotPatternDensityMap on −250/+250 regions centered on the dominant TSSs. TATA box position weight matrix (PWM) matches analyses was performed using the MotifScanScores function applied on the −35/−20 sequences centered on the dominant TSSs, using the TBP PWM provided in the SeqPattern package (derived from the JASPAR data base). Distribution of the best match for each sequence was then plotted. Sequence Logo were created using Bioconductor R package SeqLogo.

## Data availability

The datasets generated during the current study are available in different repositories: proteomic data; ProteomeXchange PRIDE database (PXD0316347 [http://www.ebi.ac.uk/pride/archive/projects/PXD016347]), RNA-seq data; Gene Expression Omnibus database (GSE140090 [https://www.ncbi.nlm.nih.gov/geo/query/acc.cgi?acc=GSE140090]”) and SLIC-CAGE data; ArrayExpress (E-MTAB-8866 [https://www.ebi.ac.uk/arrayexpress/experiments/E-MTAB-8866/]).

## Code availability

RNA-seq data was analysed using Bioconductor package DESeq2, SLIC-CAGE data was analysed using Bioconductor package CAGEr. All custom code is available upon request.

## Acknowledgements

We thank D. Singer and A. Gegonne for the gift of the *Taf7*^flox^ mouse line and H. Stunnenberg for TFIIA antibodies. We would also like to thank D. Devys for critically reading the manuscript, all members of the Tora lab for thoughtful discussions and suggestions throughout the course of the work. We are grateful to I. Kukhtevich, M. Borsos, M.E. Torres Padilla, T. Gupta, L. Casini, G. Barzaghi and A. Krebs for help in preliminary experiments. We thank C. Hérouard and M. Jung from the GenomEAST platform for library preparation and preliminary analyses, P. Eberling for peptide synthesis, F. Ruffenach for proteomic analyses, G. Duval for polyclonal antibody generation, the histology platform, the IGBMC cell culture facility and S. Falcone, M. Poirot and F. Memedov of the IGBMC animal facility for animal care taking. This work was supported by funds from CNRS, INSERM, and Strasbourg University. This study was also supported by the European Research Council (ERC) Advanced grant (ERC-2013-340551, Birtoaction) and grant ANR-10-LABX-0030-INRT (to LT) and a French State fund managed by the Agence Nationale de la Recherche under the frame program Investissements d’Avenir ANR-10-IDEX-0002-02. IB and FM acknowledge support by Wellcome Trust Senior Investigator awards (106115/Z/14/Z and 106955/Z/15/Z, respectively).

## Authors’ contribution

CY, SDV and LT designed the study; SDV and LT supervised the project; CY performed all molecular lab and mouse experiments, VH performed immunolocalizations and helped for the histological analysis of ovaries, EG generated the anti-TBPL2 polyclonal antibodies, LN carried out the proteomic analyses, KG and IB carried out preliminary analyses, PH organized the SLIC-CAGE, NC carried out SLIC-CAGE analyses, NC and BL analysed the SLIC-CAGE data, and SDV analysed the proteomic, RNA-seq and SLIC-CAGE data. FM oriented the promoter analyses. CY, FM, SDV and LT wrote the manuscript with contributions to manuscript text and figure legends from all authors. All authors gave final approval for publication.

## Competing interests

The authors declare that they have no competing interests.

## Additional Information

Supplementary information is available for this paper.

Correspondence and request for materials should be addressed to S. D. Vincent or L. Tora.

## Notes

### Competing Interest Statement

The authors have declared no competing interest.

