## Supplementary information for "TBPL2/TFIIA complex establishes the maternal transcriptome by an oocyte-specific promoter usage"

Yu *et al.*

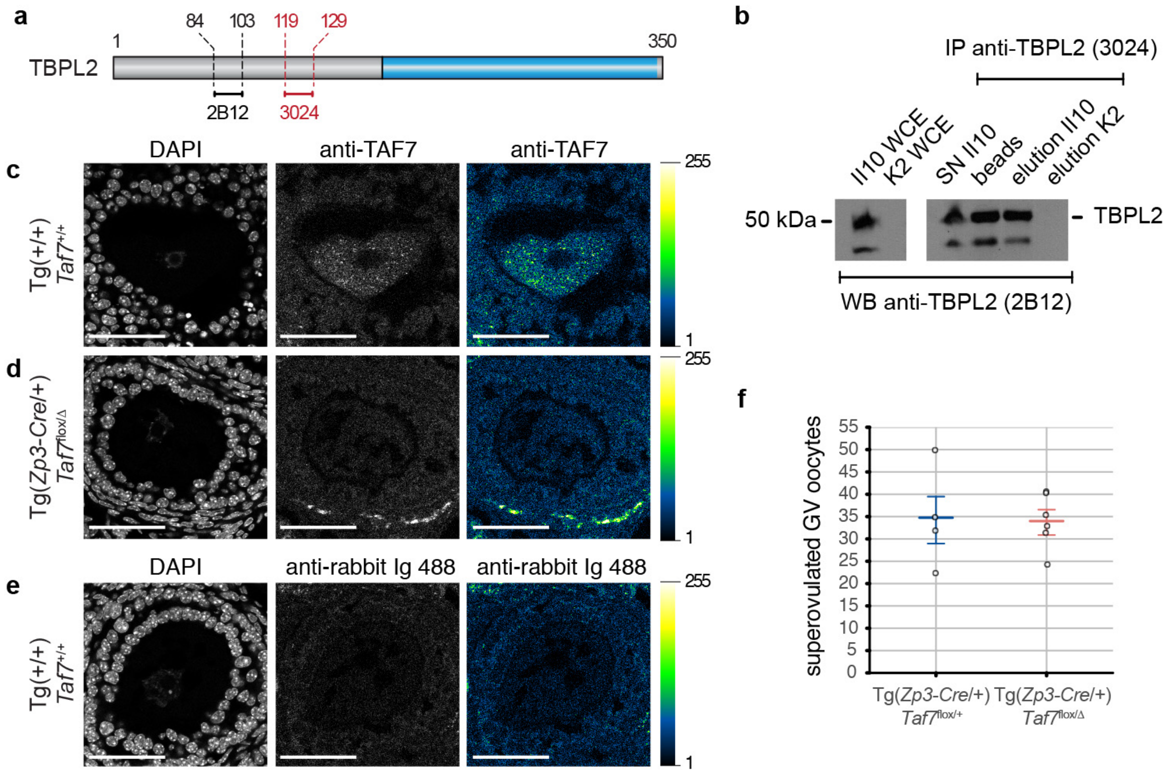


**g**

| cage n°1 | cage n°2 | cage n°3 |
|---|---|---|
| ♂ WT x 2 ♀ Tg(+/+);*Taf7*$^{flox/flox}$ | ♂ WT x 2 ♀ Tg(*Zp3-Cre*/+);*Taf7*$^{flox/\Delta}$ | ♂ WT x 2 ♀ Tg(*Zp3-Cre*/+);*Taf7*$^{flox/\Delta}$ |
| 20 | 3 | 2 |
| 11 | - | - |
| 7 | - | - |
| 9 | - | - |
| 9 | - | - |
| 11 | - | - |
| 5 | - | - |
| 72 | 3 | 2 |

**Supplementary Fig. 1** TBPL2 does not assemble in a TFIID-like complex during oocyte growth. **a** Scheme of the TBPL2 protein: the peptides used to raise the different anti-TBPL2 antibodies used are indicated (mouse monoclonal 2B12 and rabbit polyclonal 3024). **b** The anti-TBPL2 3024 polyclonal antibody (pAb) works in immunoprecipitation (IP) from whole cell extract (WCE) prepared from NIH3T3 over-expressing TBPL2 (NIH3T3-II10), as revealed by on Western blot assay, as indicated. **c-e** Representative views of immunolocalization on ovary cryosections from control (**c, e**) and oocyte-specific *Taf7* mutant (Tg(*Zp3-Cre*/+);*Taf7*$^{flox/\Delta}$ (**d**) ovaries using a purified rabbit anti-TAF7 revealed by a secondary anti-rabbit antibody coupled to Alexa 488 (**c, d**) or only the secondary anti-rabbit Ig antibody coupled to Alexa 488 (**e**). Left panels; DAPI, middle and right panels (grey scale); antibody staining in grey scale (middle panel) or Green Fire Blue LUT scale (right panel, scale on the right). Scale bars, 50 µm, analyses of 3 sections from 3 biological replicates in 2 independent experiments. **f** Number of germinal vesicle (GV) oocytes obtained after superovulation from control (Tg(*Zp3-Cre*/+);*Taf7*$^{flox}$/+, n= 4) and oocyte-specific *Taf7* mutant (Tg(*Zp3-Cre*/+);*Taf7*$^{flox/\Delta}$, n= 6) ovaries. Error bars indicate mean +/- standard error of the mean, ns; non-significant after a two-tailed Wilcoxon rank-sum test. **g** Number of pups recorded over a 7-months period from one ♂ WT x 2 ♀ Tg(+/+);*Taf7*$^{flox/flox}$ (cage n°1) and 2 ♂ WT x 2 ♀ Tg(*Zp3-Cre*/+);*Taf7*$^{flox/\Delta}$ (cages n°2 and 3) breeding cages.

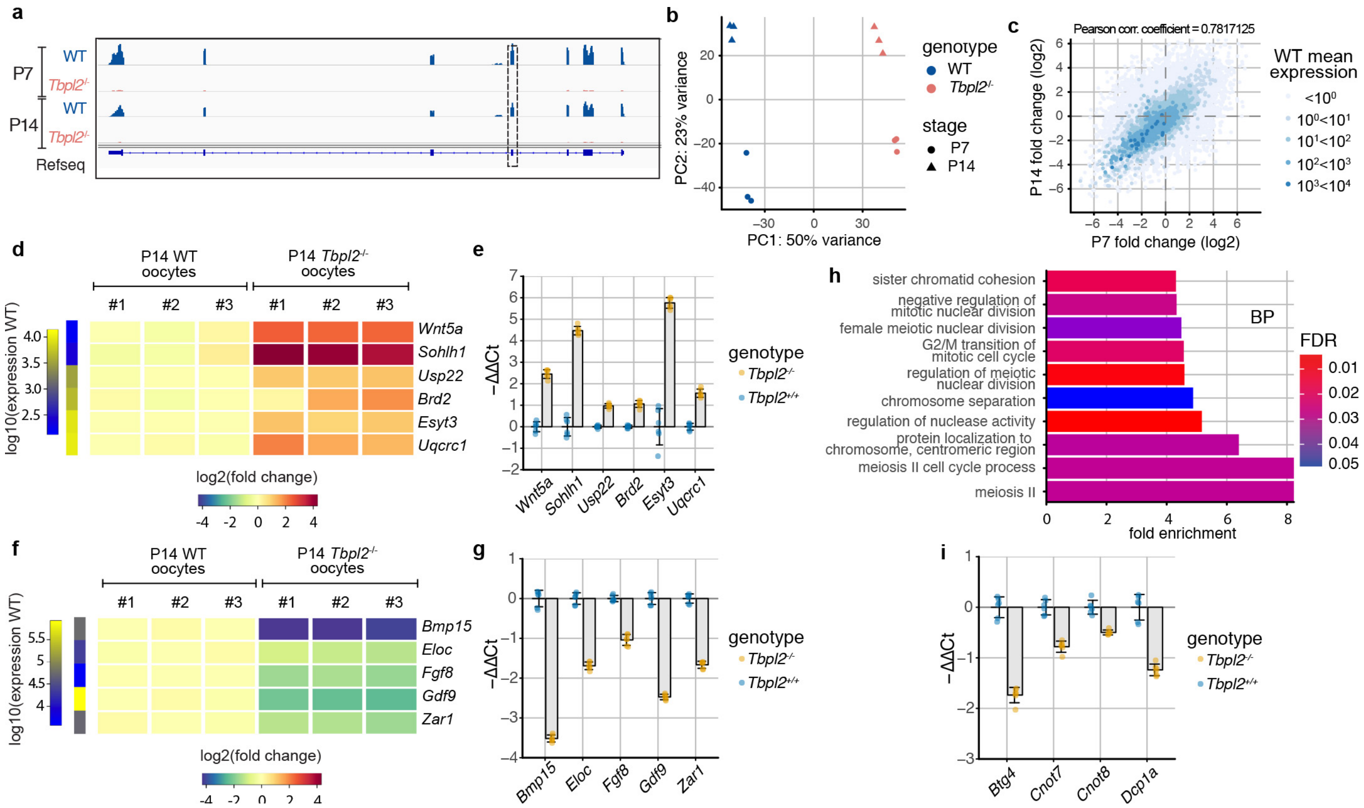


**Supplementary Fig. 2** Transcriptomic analyses of *Tbpl2*$^{-/-}$ mutant growing oocytes. **a** Normalized Integrative Genomic Viewer (IGV) snapshots for the *Tbpl2* locus. The exon 4 of *Tbpl2*, deleted in the mutant, is highlighted by a dashed box. **b** Principal component analysis after regularized log transformation of all data. **c** Post-natal day (P) 7 versus P14 fold change comparison of the color-scaled expression normalized to the median size of the transcripts in kb indicating that loss of TBPL2 has a major effect on the expression of the most abundant genes in oocytes at P7 and P14. The Pearson correlation coefficient for the transcripts expressed above 100 normalized reads is indicated on the top of the graph. **d** Heatmap of selected up-regulated candidate genes (*Brd2*, *Esyt3*, *Sohlh1*, *Uqcrc1*, *Usp22* and *Wnt5a*). Expression level in P14 WT oocytes is indicated on the first right column, the fold change for the three P14 WT and three P14 *Tbpl2*$^{-/-}$ mutant oocytes biological replicates are presented, the color legend is indicated at the bottom. **e** RT-qPCR validation of the up-regulation of the expression of *Brd2*, *Esyt3*, *Sohlh1*, *Uqcrc1*, *Usp22* and *Wnt5a* in P14 *Tbpl2*$^{-/-}$ mutant oocytes. The individual data points are indicated (biological duplicates in technical triplicates). **f** Heatmap of selected genes known to be expressed in growing oocytes (*Bmp15*, *Eloc*, *Fgf8*, *Gdf9* and *Zar1*). Expression level in P14 WT oocytes is indicated on the first right column, the fold change for the three P14 WT and three P14 *Tbpl2*$^{-/-}$ mutant oocytes biological replicates are presented, the color legend is indicated at the bottom. **g** RT-qPCR validation of the down-regulation of the expression of *Bmp15*, *Eloc*, *Fgf8*, *Gdf9* and *Zar1* in P14 *Tbpl2*$^{-/-}$ mutant oocytes (biological duplicates in technical triplicates). **h** Down-regulated genes GO category analyses for biological processes (BP). The top ten most enriched significant GO categories for a FDR $\leq 0.05$ are represented. **i** RT-qPCR validation of down-regulation of the expression of *Btg4*, *Cnot7*, *Cnot8* and *Dcp1a* in P14 *Tbpl2*$^{-/-}$ mutant oocytes. The individual data points are indicated on the figure (biological duplicates in technical triplicates). Error bars indicate mean +/- standard deviation in (**e, g, i**).

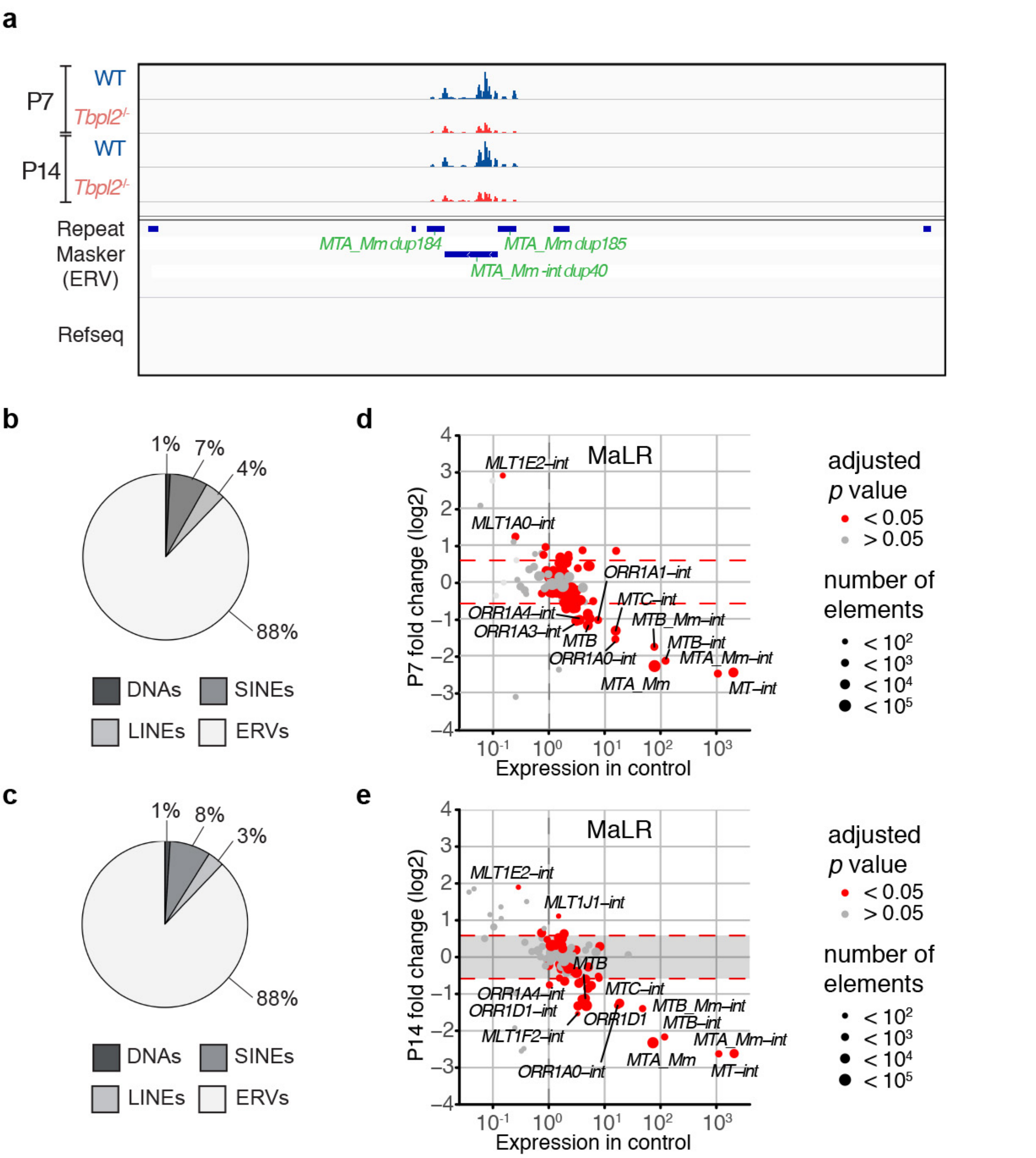


**Supplementary Fig. 3** Transcriptomic analyses of Endogenous Retroviral Elements (ERVs) of *Tbpl2*$^{-/-}$ mutant growing oocytes. **a** Normalized Integrative Genomic Viewer (IGV) view of a region devoid of Refseq annotated genes but containing multiple mammalian apparent LTR retrotransposons (MaLR) endogenous retroviral elements (ERV) annotated in the RepMasker data base at post-natal day (P) 7 and P14. Reads accumulation corresponding to the *MTA_Mm_dup184*, *MTA_Mm-int_dup40* and *MTA_Mm_dup185* elements is diminished in P7 and P14 *Tbpl2*$^{-/-}$ mutant oocytes. **b**, **c** Distribution of the expression of the major classes of repeat elements in WT P7 (**b**) and P14 (**c**) oocytes indicating that the ERV retrotransposon class is the most abundantly expressed. **d**, **e** Differential expression of the ERV MaLR family members indicating that the *MTA_Mm*, *MTA_Mm-int* and *MT-int* elements are the most severely affected in *Tbpl2*$^{-/-}$ mutant oocytes at P7 (**d**) and P14 (**e**), adjusted *p* value after two-sided Wald test and Benjamini-Hochberg correction for multiple comparison.

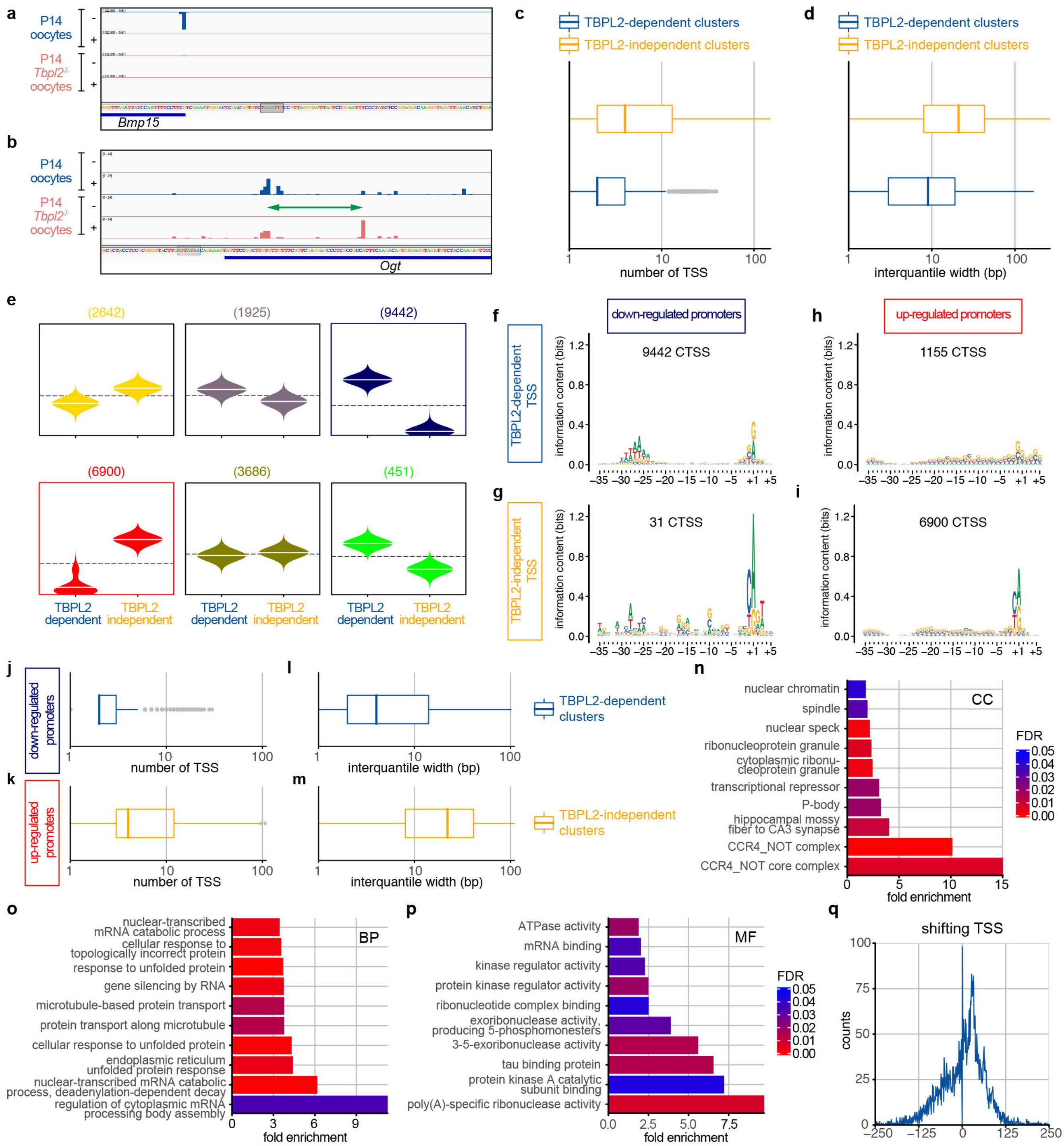


**Supplementary Fig. 4** Core promoter regions of TBPL2 specific transcription units in P14 oocyte are enriched in TATA-like elements and are sharp. **a, b** Integrative Genomic Viewer (IGV) snapshots of post-natal day (P) 14 wild-type (WT) (blue) and *Tbpl2*$^{-/-}$ mutant (salmon) oocytes SLIC-CAGE data. Some TSSs of genes expressed in WT oocytes and downregulated in *Tbpl2*$^{-/-}$ mutants display very sharp TSS patterns (**a**). TSSs from genes expressed both in WT and *Tbpl2*$^{-/-}$ mutant oocytes can display a shifting TSS pattern (green double arrow). (**b**). Putative TATA-like elements are highlighted in grey. **c, d** Promoter architecture analyses of TBPL2-dependent (blue, n=32515) and TBPL2-independent (orange, n=23932) TSS clusters: distribution of TSSs number (**c**) and of cluster interquantile width (**d**). **e** Expression profiling of TBPL2-dependent and TBPL2-independent consensus TSS clusters using self-organizing map (SOM) clustering. Numbers in brackets indicate the number of consensus clusters for each SOM group. The two selected SOM groups are boxed in blue (down-regulated promoters) and red (up-regulated promoters). **f-i** Sequence logo analyses of the -35/+5 sequence region of the dominant TSS (+1) for the TBPL2-dependent (**f, h**) and TBPL2-independent (**h, i**) cluster TSSs (CTSS) for the down-regulated promoters (**f, g**) and up-regulated promoters (**h, i**) SOM groups. The number of TSS clusters present in each group is indicated on the top of each panel. **j-m** Promoter architecture analyses of TBPL2-dependent TSS clusters of the down-regulated promoters (**j, l**, n=9442) and of TBPL2-independent TSS clusters of the up-regulated promoters (**k, m**, n=6900) groups: distribution of TSSs number (**j, k**) and of cluster interquantile width (**l, m**) of the respective TSS clusters. **n-p** Down-regulated promoters GO category analyses for cellular component (CC, **n**), biological process (BP, **o**) and molecular function (MF, **p**). The top ten most enriched significant GO categories for a FDR ≤ 0.05 are represented. **q** Distribution of the distance between the dominant TSS of the TBPL2-dependent versus TBPL2-independent shifting promoters (6433 shifting TSS clusters). The distance width has been restricted to -250 and +250 since it contains >95% of the information. Boxplots in (**c, d, j-m**) show the $5^{th}$, $25^{th}$, $50^{th}$, $75^{th}$, and $95^{th}$ percentiles and center line is the median, outliers are indicated in grey dots.

**Supplementary Table 1 :** list of antibodies

| primary antibody | name | type | source | reference |
|---|---|---|---|---|
| anti-TBPL2 | 3024 | rabbit polyclonal | IGBMC | this study |
| anti-TBPL2 | 2B12 | mouse monoclonal | IGBMC | [1] |
| anti-TAF7 | 3475 | rabbit polyclonal | IGBMC | [2] |
| anti-TFIIA-alpha | | rabbit polyclonal | gift from HG Stunnenberg | [3] |
| anti-TBP | 3TF1-3G3 | mouse monoclonal | IGBMC | [4] |
| anti-TAF6 | 25TA2G7 | mouse monoclonal | IGBMC | [5] |
| anti-TAF10 | 6TA2B11 | mouse monoclonal | IGBMC | [6] |
| anti-GST | 15TF21D10 | mouse monoclonal | IGBMC | [7] |

| secondary antibody | supplier | reference | lot |
|---|---|---|---|
| Alexa Fluor 488 goat anti-rabbit Ig | Invitrogen | #A-11108 | 1829924 |
| Peroxidase coupled Goat Anti-Mouse IgG | Jackson ImmunoResearch | #115-036-071 | 132326 |
| Peroxidase coupled Goat Anti-Rabbit IgG | Jackson ImmunoResearch | #115-035-144 | 132676 |

**Supplementary Table 2 :** list of primers

| gene | primer | sequence | reference |
|---|---|---|---|
| *18rRNA* | forward | GTAACCCGTTGAACCCCATT | doi:10.1038/onc.2011.12 |
| *18rRNA* | reverse | CCATCCAATCGGTAGTAGCG | doi:10.1038/onc.2011.12 |
| *Bmp15* | forward | AAGGGAGAACCGCACGATTG | doi:10.1038/srep23972 |
| *Bmp15* | reverse | TGCTTGGTCCGGCATTTAGG | doi:10.1038/srep23972 |
| *Brd2* | forward | AACAGCCATAAGAAGGGGGC | |
| *Brd2* | reverse | AGGGGTGGTAGTATCCGCTT | |
| *Btg4* | forward | TGAAAAAGCATGAGAAACTGAGTAC | doi:10.1038/nsmb.3204 |
| *Btg4* | reverse | CCCATCTACCTTTAAAAGAAGCAA | doi:10.1038/nsmb.3204 |
| *Cnot7* | forward | GGTGGATTACAGGAAGTTGCTG | doi:10.1038/nsmb.3204 |
| *Cnot7* | reverse | GGATGAGCCAGAACCAAGG | doi:10.1038/nsmb.3204 |
| *Cnot8* | forward | CATTCTGCTTCTGGGCCAC | doi:10.1038/nsmb.3204 |
| *Cnot8* | reverse | CTTTTGGAAGCAGAGTCCGA | doi:10.1038/nsmb.3204 |
| *Dcp1a* | forward | GACTGTCATCGCATAGCA | |
| *Dcp1a* | reverse | CCTCTCATACTCATCCTTGG | |
| *Eloc* | forward | ATCTTCTGATGGCCATGAATTT | doi:10.1038/nsmb.3204 |
| *Eloc* | reverse | CCTTGTAGGTAAAATACATGCACAC | doi:10.1038/nsmb.3204 |
| *Esyt3* | forward | GGTTTGCCCTGAATGACACG | |
| *Esyt3* | reverse | CTGGCCTTATTCGGCAGGTT | |
| *Fgf8* | forward | CTTTTGGAAGCAGAGTCCGA | doi:10.1038/nsmb.3204 |
| *Fgf8* | reverse | CCATGTACCAGCCCTCGTAC | doi:10.1038/nsmb.3204 |
| *Gdf9* | forward | GATGGGACTGACAGGTCTGG | doi:10.1038/srep23972 |
| *Gdf9* | reverse | CAGCGGTCCTGTCACCTG | doi:10.1038/srep23972 |
| *Sohlh1* | forward | CTCAGGGTCTCCGATGAAGC | |
| *Sohlh1* | reverse | CTCCCGAGGCAAGTGACATT | |
| *Uqcrc1* | forward | ATGCTGCGTGACATTTGCTC | |
| *Uqcrc1* | reverse | CGGTTGTAGTCTGGGAGCTG | |
| *Usp22* | forward | AGGGCAGTGTGGTTAATGGG | |
| *Usp22* | reverse | ATGGCAACCGCTACACTTGA | |
| *Wnt5a* | forward | GCAGGACCTGGTCTACATCG | |
| *Wnt5a* | reverse | GTCCATCCCCTCTGAGGTCT | |
| *Zar1* | forward | AGAGCGCCTATGTGTGGTGT | doi:10.1038/nsmb.3204 |
| *Zar1* | reverse | TCTCCCACACAAGTCTTGCC | doi:10.1038/nsmb.3204 |
